# Regenerative epithelial spike cells instruct immune niche homeostasis in zebrafish fin epidermis

**DOI:** 10.64898/2026.11.05.764666

**Authors:** Siyang Cao, Margaret K. Keating, Julia O’Hagan, Anjelica Rodriguez-Parks, Yiran Hou, Tanner F Robertson, Anna Huttenlocher, Junsu Kang

## Abstract

Epithelia comprise diverse specialized epithelial cells that interact with immune cells, yet how these interactions sustain immune cell pools remain poorly understood. Here, we integrated single-cell RNA sequencing (scRNA-seq) analysis with transgenic assays in zebrafish and discovered a previously unrecognized epithelial population marked by *advillin* (*avil)*. We identified epithelial cells exhibiting unique morphology and localization, characterized by an apical spike projecting outward from the body - which we term spike cells. These cells exhibit features reminiscent of mammalian tuft cells. Notably, *avil*^+^ spike cells exhibit close spatial association with a subset of *mpeg1*^+^ immune cells. Following fin amputation, spike cells are replenished *de novo* from precursors and undergo dynamic changes in density, morphology, and orientation, and their regeneration depends on Alk signaling. Moreover, spike cells dynamically interact with *mpeg1^+^* immune cells to maintain the immune cell population. Together, our findings identify a unique epithelial population in zebrafish fins and define their roles in maintaining immune cell niche dynamics.

## Introduction

Epithelia cover the external surfaces of the body and line internal organs and cavities, serving as protective barriers against pathogens and environmental hazards^1^. Beyond their barrier function, different epithelial tissues contain specialized cell populations that perform diverse physiological roles. Goblet cells secrete mucus to lubricate surfaces and trap harmful substances and pathogens^2^. Chemosensory epithelial cells detect chemical cues and contribute to local tissue responses^3, 4, 5^. Somatosensory cells, such as Merkel cells and Meissner’s corpuscles, enable the detection of diverse stimuli and transmission of signals to the nervous system^6, 7^. Club cells in the skin of fish species release alarm substances to warn conspecifics of potential threats^8^. Maintaining appropriate pools of these specialized epithelial cells is critical for preserving tissue function and ensuring effective responses to environmental cues.

Zebrafish inhabit aquatic environments, which impose unique environmental challenges compared with those of terrestrial mammals. While keratinocytes are the predominant cell type in zebrafish skin^9^, the epidermis also contains specialized cells, such as ionocytes, which maintain ionic homeostasis^10, 11^, and goblet cells, which produce the mucus layer^12, 13^. During early embryogenesis, the zebrafish epidermis consists of two keratinocyte layers interspersed with specialized epidermal cells, such as ionocytes and goblet cells^14^. During postembryonic development, as zebrafish reach approximately 5.0–6.0 mm standard length (SL), additional cell layers are added through a process known as epidermal stratification^15, 16^. At the onset of stratification, flattened basal cells undergo a morphological shift to a more rounded shape. In addition, two distinct populations contribute to epidermal stratification: slowly proliferative basal epidermal stem cells with high p63 expression and rapidly proliferative progenitor cells with lower p63 expression^15^.

These populations collectively give rise to a mature epidermis composed of three to four layers. In adult zebrafish, the epidermis exhibits remarkable regenerative capacity, efficiently replenishing lost cells following fin amputation^17, 18^. However, the composition of specialized epidermal cell populations in the mature epidermis and the mechanisms governing their regeneration remain largely unknown.

Tuft cells - also known as brush cells, microvillous cells, and caveolated cells - are solitary chemosensory epithelial cells that serve as sentinels in mucosal tissues^19, 20, 21^. Initial studies in the mammalian intestine revealed that tuft cells detect parasite-derived stimuli and relay these signals to activate group 2 innate lymphoid cells (ILC2s), highlighting their immune-related functions^5, 22, 23^. Recent research has highlighted the heterogeneity of tuft cells across tissue types and anatomical locations^24, 25, 26^. Furthermore, tuft cells are increasingly recognized for their multifaceted functions, including sensing microorganisms, monitoring environmental changes, mediating cholinergic signaling, and contributing to tumorigenesis^5, 27, 28, 29, 30, 31, 32, 33, 34^. In zebrafish, the presence of tuft cells has been suggested by recent single-cell RNA sequencing (scRNA-seq) studies^35, 36, 37, 38^. However, the morphological features and dynamic cellular characteristics of tuft cells in zebrafish remain largely unexplored.

In this work, we report a previously uncharacterized epithelial cell population in zebrafish skin. Within the fin epidermis, we identified tuft cell-like epithelial cells that display a distinctive spike-shaped apical morphology and express tuft cell markers. These spike cells are located in close proximity to *mpeg1*^+^ immune cells in the fin epidermis. Following amputation, spike cells regenerate dynamically from epidermal precursor cells. Moreover, the presence of spike cells is required to maintain the local immune cell population in fin tissues. Collectively, our findings provide evidence for a unique epithelial cell population analogous to mammalian tuft cells and establish a model for further investigation of their function in regulating immune cell niches.

## Results

### scRNA-seq analysis identifies diverse epithelial clusters in regenerating fin tissues

To investigate the diverse and dynamic cellular states associated with adult zebrafish caudal fin regeneration, we analyzed two published scRNA-seq datasets (**Fig. 1a**). The first dataset consists of cells dissociated from regenerating adult fins at 1 day post-amputation (1 dpa), referred to here as O1dpa (only 1 dpa samples)^39^. The second dataset contains cells dissociated from uninjured fins and regenerating adult fins collected at 1, 2, and 4 dpa, referred to here as M-fins (multiple time points)^40^. To avoid batch effects, we analyzed each dataset independently and compared clusters based on shared or distinct gene expression patterns. After quality-control filtering, 12,503 and 23,702 cells remained for O1dpa and M-fins, respectively.

**Figure 1.**
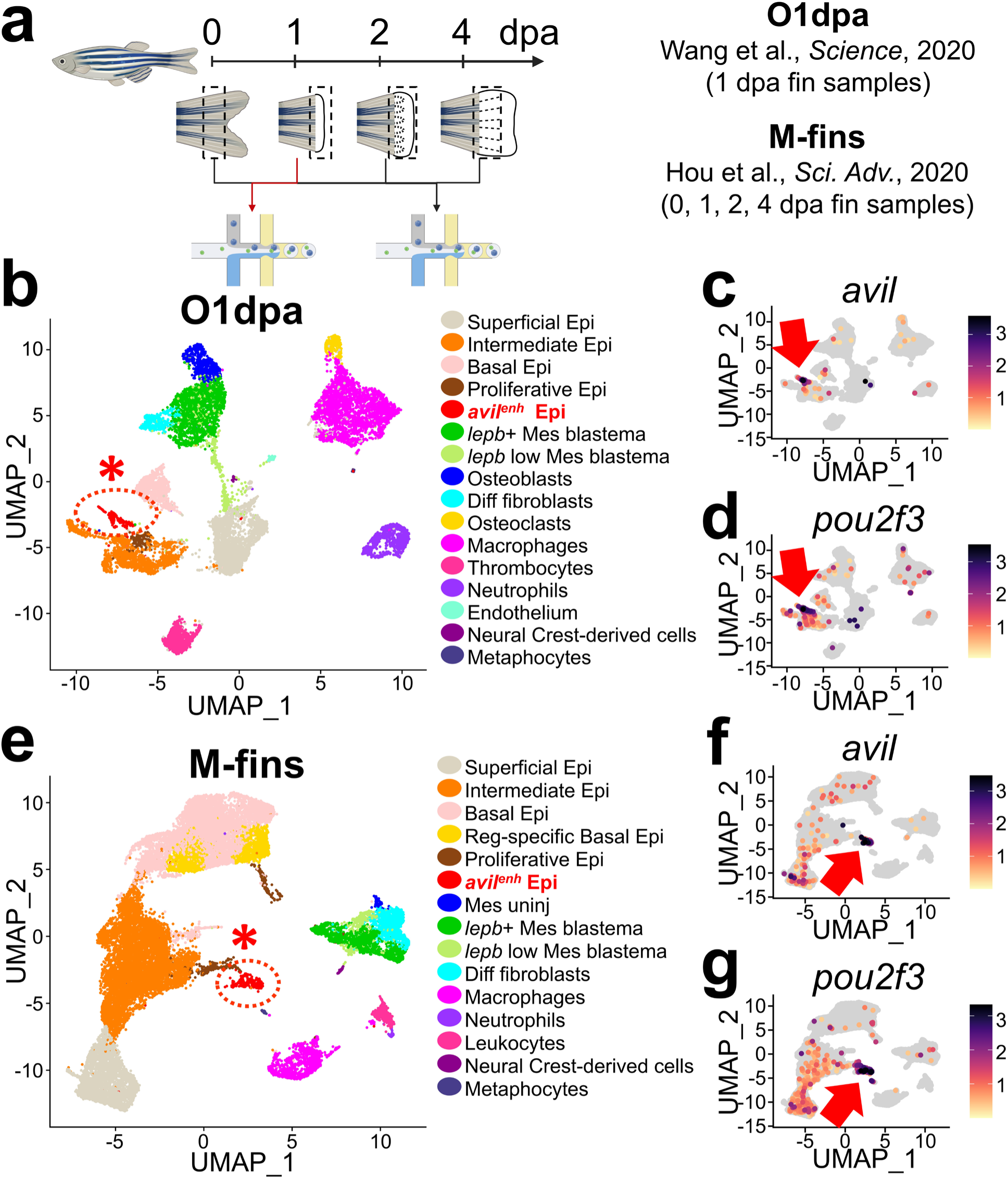
scRNA-seq analyses identify *avil*^+^ tuft cell-like epithelial cells. (**a**) A schematic of scRNA-seq analyses with fin tissues. O1dpa and M-fins datasets are obtained from PRJNA559885 and GSE137971, respectively. (**b**) Uniform Manifold Approximation and Projection (UMAP) of fin tissues collected at 1 day post-amputation (1 dpa). (**c, d**) Expressions of *avil* (**c**) and *pou2f3* (**d**) in O1dpa dataset. (**e**) UMAP of uninjured and regenerating fin tissues. (**f, g**) Expressions of *avil* (**f**) and *pou2f3* (**g**) in M-fins dataset. The asterisks indicate *avil*-enriched epidermis (*avil^enh^* Epi) population.

Unbiased transcriptomic analyses identified 16 and 15 cell clusters in O1dpa and M-fins, respectively. Epithelial cells marked by *epcam* and *cdh1*^40^, mesenchymal cells or fibroblasts marked by *msx1b* and *msx3*^41, 42^, macrophages marked by *mpeg1.1* and *mfap4* ^43, 44^, neutrophils marked by *mpx* and *lyz*^45, 46^, neural crest-derived cells marked by *foxd3* and *sox10*^47, 48^ and metaphocytes marked by *spic*^49^ were identified in both datasets (**Fig. 1b, c** and **S1a, b**). Leukocytes marked by *coro1a* and *cxcr4b* ^50, 51^ were observed in M-fins, whereas osteoclasts marked by *acp5a/trap* and *spp1*^52^, endothelial cells marked by *kdrl, cdh5*, and *ecscr* ^53, 54, 55, 56^ and thrombocytes marked by *gp1bb* and *hbaa2* ^57^ were detected only in O1dpa (**Fig. 1b, c** and **S1a, b**), presumably due to technical differences in sample preparation.

We focused on epithelial cells, the most abundant cell population in both datasets. Consistent with previous reports^35, 40, 58^, our scRNA-seq analysis identified epithelial populations corresponding to three major epithelial layers: superficial, intermediate, and basal epithelium, characterized by distinct expression patterns of *claudin* genes (**Fig. S1a, b**). *cldnb* was enriched in both superficial and intermediate epithelial layers. Whereas *cldna* was strongly expressed in intermediate epidermis, *cldni* was prominent in the basal epidermis. The transcription factor p63, a master regulator of epidermal development ^15, 59^, was enriched in intermediate and basal epithelial layers but was rarely detectable in superficial epithelial cells. We also identified a proliferative *p63*^+^ epithelial cluster expressing genes related to cell cycle progression and proliferation, such as *cdk1*, *anln, and plk1* (**Fig. S1a, b**). Interestingly, our scRNA-seq analysis identified a specialized epithelial cell cluster with low p63 expression (**Fig. 1b, e and S1a, b**). Gene ontology (GO) analysis^60^ of genes enriched in this cluster, using thresholds of adjusted P value < 0.05 and log2FC > 1, highlighted terms related to epithelial and immune-associated functions (**Fig. S1c**). We found that *advillin* (*avil*) was highly enriched in this specialized epithelial cluster (**Fig. 1c, g**). Recent studies have reported *avil* as a marker of mammalian tuft cells, a specialized epithelial cell type involved in pathogen sensing and intestinal immunity^24, 61, 62^. Consistent with these findings, several tuft cell markers, including *pou2f3*, *gng13a*, *tuba1a*, and *rgs13*^22, 62, 63, 64, 65^, were highly expressed in this cluster (**Fig. 1d, h, and S1a, b**). Collectively, our scRNA-seq analysis uncovered diverse epithelial cell populations in zebrafish fin tissues, including a tuft cell-like epithelial population.

### *avil*^+^ cells represent a specialized spike-shaped epithelial cell population

To visualize *avil-*expressing cells, we generated a transgenic zebrafish reporter line using *avil* regulatory sequences, including 443-bp genomic sequence upstream of the start codon and 512-bp downstream genomic sequence located within the adjacent intron. *avil:memb-mCherry* (*avil*:*m-mCherry*) expresses mCherry fused to the N-terminal membrane-targeting sequence of zebrafish Gap43^66, 67^. The resulting fluorescent signal is enriched at the plasma membrane but is also detectable in intracellular regions, while being excluded from the nucleus. By 5 days post-fertilization (dpf), *avil:m-mCherry* labeled cells in the olfactory region and taste buds (**Fig. S2a**). By 14 dpf, *avil:m-mCherry* expression emerged in the trunk epidermis in addition to the olfactory region (**Fig. S2b**). In adult zebrafish, *avil^+^* cells were sparsely distributed throughout the skin epidermis (**Fig. S2c**). In caudal fins, *avil*^+^ cells appeared as individual spherical cells throughout the fin epidermis and were more densely distributed in intra-fin ray regions than inter-fin ray regions (**Fig. 2A** and **S2d)**. Using an *avil* HCR probe, we confirmed that endogenous *avil*-expressing cells correspond to *avil:m-mCherry-*expressing cells (**Fig. S2e**).

**Figure 2.**
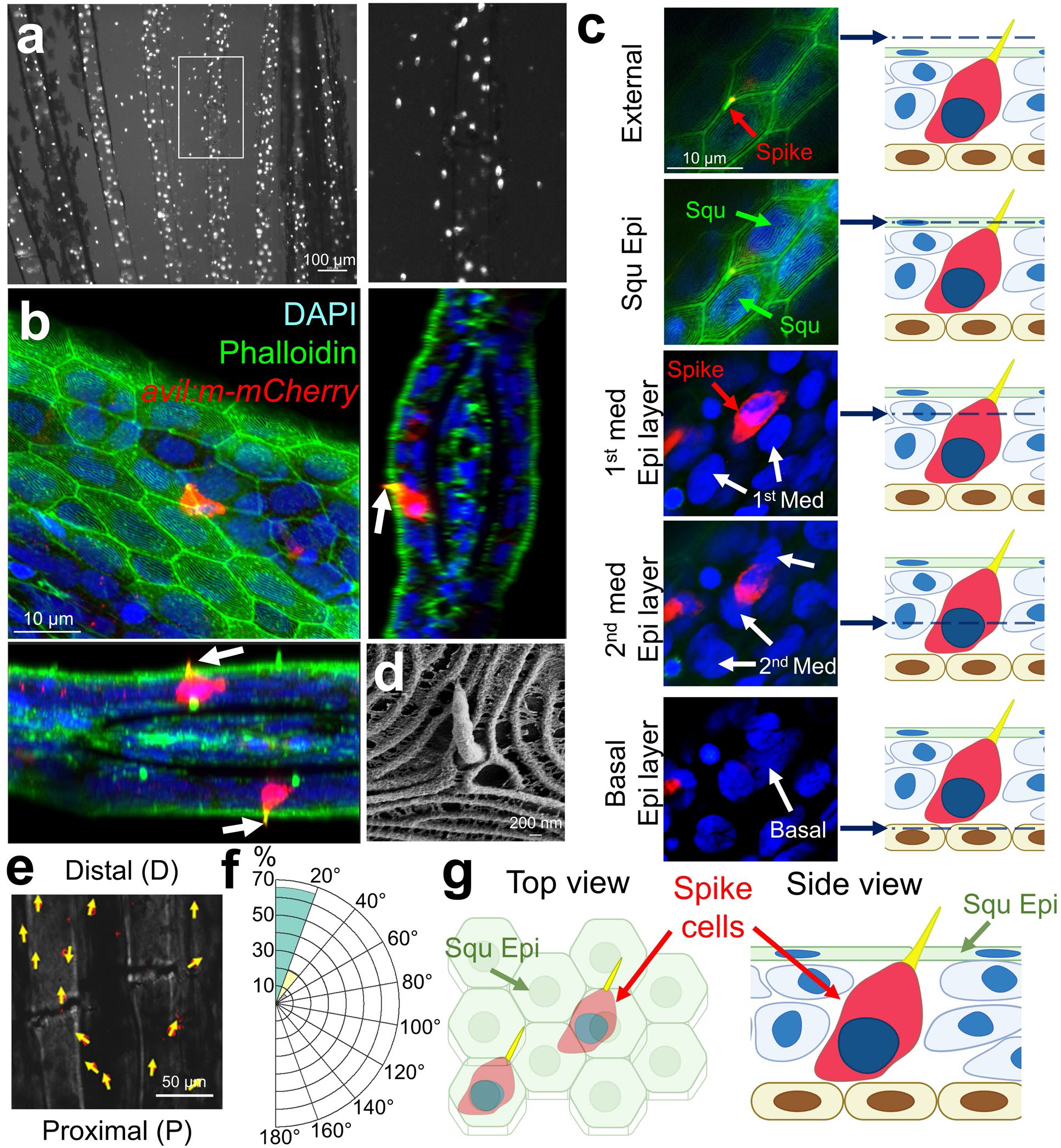
*avil*^+^ spike cells display unique morphology and anatomical location in adult zebrafish fin epidermis. (**a**) Whole-mount image of *avil:m-mCherry* displays scattered *avil*^+^ cells distributed throughout the adult fin tissues. (**b**) Confocal whole-mount image of uninjured *avil:m-mCherry* adult fins stained with anti-mCherry antibody and Phalloidin. Squamous epidermal cells (Squ Epi) are characterized by microridge structures on their apical surface. Orthogonal x-z and y-z projection views are shown at the bottom and right, respectively. Arrows indicate F-actin-enriched spike structures. (**c**) Serial confocal sections from the external to the basal cell layer, with corresponding z-axis positions indicated in the schematic on the right. Med: Intermediate epidermal cells. (**d**) Scanning electron microscopy (SEM) image of the spike cell in fin tissues. (**e**) Representative whole-mount image of *avil:m-mCherry* signals overlaid with bright-field image. Arrows indicate the orientation of spike cells from the proximal (P) to the distal tip (D). (**f**) A rose plot depicting the orientation of spike cells, with 0° and 180° representing D and P directions, respectively (n = 384 cells from 6 fish). (**g**) Schematic of top and side views of uninjured fin tissues highlighting spike cells (Red). Data are presented with mean ± SEM. Green: Squ Epi. Yellow: F-actin-enriched spike structure. Blue: Intermediate (Med) epidermal cells. Light brown: Basal epidermal layer. Scale bars: **a,** 100 µm; **b, c,** 10 µm; **d,** 200 nm; **e,** 50 µm.

Specialized epithelial cells, including tuft cells, often display distinctive morphological features ^68^. To examine the morphology of *avil*^+^ cells, we performed whole-mount confocal microscopy on uninjured fins stained with anti-mCherry antibody and phalloidin to visualize F-actin. Whole-mount confocal imaging revealed that *avil*^+^ cells possess a unique morphology and localization (**Fig. 2b, c**). *avil^+^* cells are positioned directly beneath the superficial squamous epithelial layer, while extending an elongated apical spike outward from the body (**Fig. 2b, c, S2f, and Supplementary Movie 1**). The spike tip is enriched in actin filaments (F-actin), indicating a specialized apical structure (Arrows in **Fig. 2b** and **S2f**). Scanning electron microscopy (SEM) further confirmed the presence of this distinctive apical spike in the caudal fin epidermis (**Fig. 2d**). Based on this prominent apical projection, we designate these *avil⁺* cells as spike cells. We further confirmed that *avil*^+^ spike cells differ from other specialized epithelial cells, including cells of the lateral line system and Merkel cells (**Fig. S2g-i**). Aside from the spike, individual *avil*^+^ spike cells span nearly the full thickness of the stratified epithelium, extending from the superficial squamous layer to just above the basal layer (**Fig. 2b, c**), indicating that *avil*⁺ spike cells are markedly large epithelial cells.

To determine developmental appearance of *avil*^+^ spike cells, we analyzed *avil:lifeact-mNeonGreen* (*avil:l-Green*) transgenic fish that express an F-actin-binding green fluorescence reporter in *avil*-expressing cells. By 5 dpf, spike-shaped *avil:l-Green*^+^ cells were detectable primarily in craniofacial sensory regions (**Fig. S2j**). By 4.5 mm SL, *avil:l-Green*⁺ cells became detectable in the trunk epidermis (**Fig. S2k**), with exhibiting the spike morphology. By 6.0 mm SL, *avil:l-Green*⁺ spike cells were markedly more abundant and broadly distributed throughout the trunk epidermis (**Fig. S2l**). As epidermal stratification begins at approximately 5.0 – 6.0 mm SL^15, 16^, these observations suggest that the robust expansion of spike cells coincides with epidermal stratification. *avil*⁺ spike cells were observed in both the trunk skin and caudal fin epidermis (**Fig. 2a, S2c** and **S2l**), demonstrating that this cell population is not restricted to the fins.

Epithelial cells often exhibit defined orientation and polarity at the tissue and cellular levels. Multicellular epithelial structures, such as fish scales and bird feathers, typically align along the anterior-posterior axis^69, 70, 71^. Individual epithelial cells can also display coordinated cellular polarity^72, 73^. For instance, *Drosophila* trichomes and vertebrate hair cells in the inner ears exhibit highly organized alignment. Because the spike-shaped morphology of *avil*^+^ cells suggests directional polarity, we examined whether spike cells exhibit a consistent orientation throughout the fin. Notably, we found that the majority of spike cells were oriented along the proximal-to-distal (P-to-D) axis. Specifically, 65.9% and 20.6% of spikes pointed distally within 0 - 20° and 20 - 40°, respectively (n = 384 cells from 6 fish; **Fig. 2e, f and S3**). Although a small proportion of spikes (4.2%) were oriented toward the distal-to-proximal direction (90–180°), the overall distribution revealed a strong preference for P-to-D orientation. Overall, these findings identify *avil*^+^ spike cells as a uniquely shaped and oriented epithelial cell population with distinct spatial organization in the zebrafish fin epidermis (**Fig. 2g**).

### *avil⁺* spike cells closely associate with *mpeg1^+^* immune cells

A subset of specialized epithelial cells functions as immune sentinels that modulate local immune responses^19, 20^. In mammals, tuft cells are chemosensory epithelial cells that orchestrate type 2 immune responses by activating ILC2s and promoting downstream immune cascades ^5, 22, 23, 29, 31^. Given that *avil*⁺ spike cells share morphological and transcriptional similarities with mammalian tuft cells, we hypothesized that they interact with immune cell populations. To test this, we crossed the *avil:m-mCherry* reporter line with *mpeg1:EGFP,* a reporter that labels multiple immune cell populations in the skin epidermis, including macrophage-lineage cells, Langerhans cells, and metaphocytes ^43, 49, 74, 75, 76^, and examined their spatial relationship in the fin epidermis. Notably, we observed that *avil*⁺ spike cells were frequently positioned in close proximity to *mpeg1^+^* cells (**Fig. 3a**). Confocal image analysis revealed that 36.9% of *avil*⁺ spike cells were likely in direct contact with *mpeg1 ^+^* cells in uninjured fin tissues (**Fig. 3b**). To determine whether this association was preferential to spike cells, we compared *mpeg1*⁺ cell contacts with cortical F-actin-enriched cells, presumably Merkel cells - another specialized epidermal cell population. A significantly higher proportion of *avil*⁺ spike cells contacted *mpeg1^+^* cells than did cortical F-actin-enriched cells, supporting a preferential association between spike cells and *mpeg1^+^* immune cells (**Fig. S4**).

**Figure 3.**
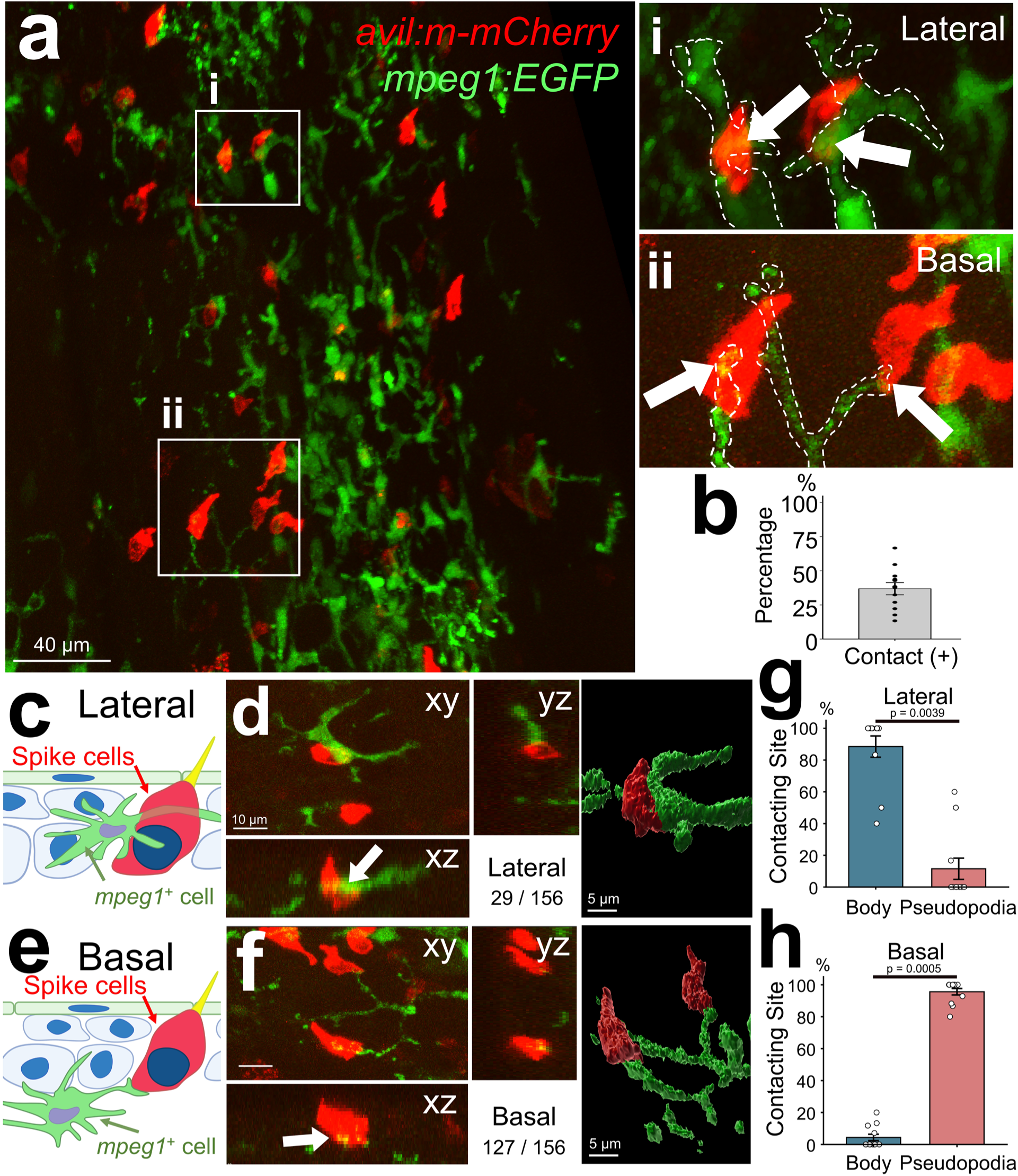
*avil*⁺ spike cells exhibit close spatial association with *mpeg1*^+^ immune cells. (**a**) Representative whole-mount confocal image of uninjured fins for *avil:m-mCherry* (red) and *mpeg1:EGFP* (green). Boxed regions (i, ii) indicate areas shown at higher magnification. (**b**) Quantification of the percentage of *avil*⁺ spike cells in contact with *mpeg1^+^* cells (n=12 fin rays from 4 fins). (**c**) Schematic of lateral contact, in which the cell body of an *mpeg1⁺* immune cell contacts the lateral membrane of an *avil*⁺ spike cell. (**d**) Three-dimensional reconstructions and orthogonal (XY and XZ) views of a representative lateral contact. (**e**) Schematic of basal contact, in which a pseudopodial process of an *mpeg1*⁺ immune cell contacts the basal surface of an *avil*⁺ spike cell. (**f**) Three-dimensional reconstructions and orthogonal views of a representative basal contact. Quantification revealed that basal contacts (n = 127) were substantially more frequent than lateral contacts (n = 29) across 12 fin-ray regions from 4 fish. (**g, h**) Quantification of the proportions of cell-body-mediated and pseudopodia-mediated contacts among basal (**g**) and lateral (**h**) contacts, respectively. Scale bars: **a,** 40 µm; **d, f,** 10 µm for orthogonal views and 5 µm for three-dimensional reconstructions.

Using three-dimensional (3D) reconstructions, surface renderings, and orthogonal (xz and yz) views, we identified two distinct modes of contact based on the location of the contact site on spike cells: lateral and basal (**Fig. 3c-f** and **Supplementary Movie 2, 3**). Quantitative analysis revealed that basal contacts were markedly more frequent than lateral contacts (81.4% and 18.6% for basal and lateral contact, respectively; n=156). In lateral contacts, the cell bodies of *mpeg1*⁺ cells mainly contacted to the lateral side of *avil*⁺ spike cells, whereas in basal contacts, the basal surface of *avil*⁺ spike cells was typically contacted by the tips of pseudopodial processes extending from *mpeg1*⁺ cells (**Fig. 3g, h**). Together, these results demonstrate that spike cells establish direct physical interactions with immune cells, suggesting a potential role in local immune modulation.

### *avil*^+^ cells robustly regenerate upon fin amputation

Zebrafish caudal fins regenerate robustly after amputation and with high fidelity^77, 78^, providing a useful model for studying the cellular and molecular mechanisms underlying the regeneration of specialized epithelial cell types. To determine whether spike cells are regenerative, we conducted fin amputation and quantified *avil^+^* cell in uninjured and regenerating fins. The number of *avil^+^* cells reached its lowest level at 2 dpa and increased thereafter, suggesting that recovery begins around this time point. By 41 dpa, spike cells were restored to levels comparable to those in uninjured fins, indicating their successful regeneration (**Fig. 4a** and **S3a**).

**Figure 4.**
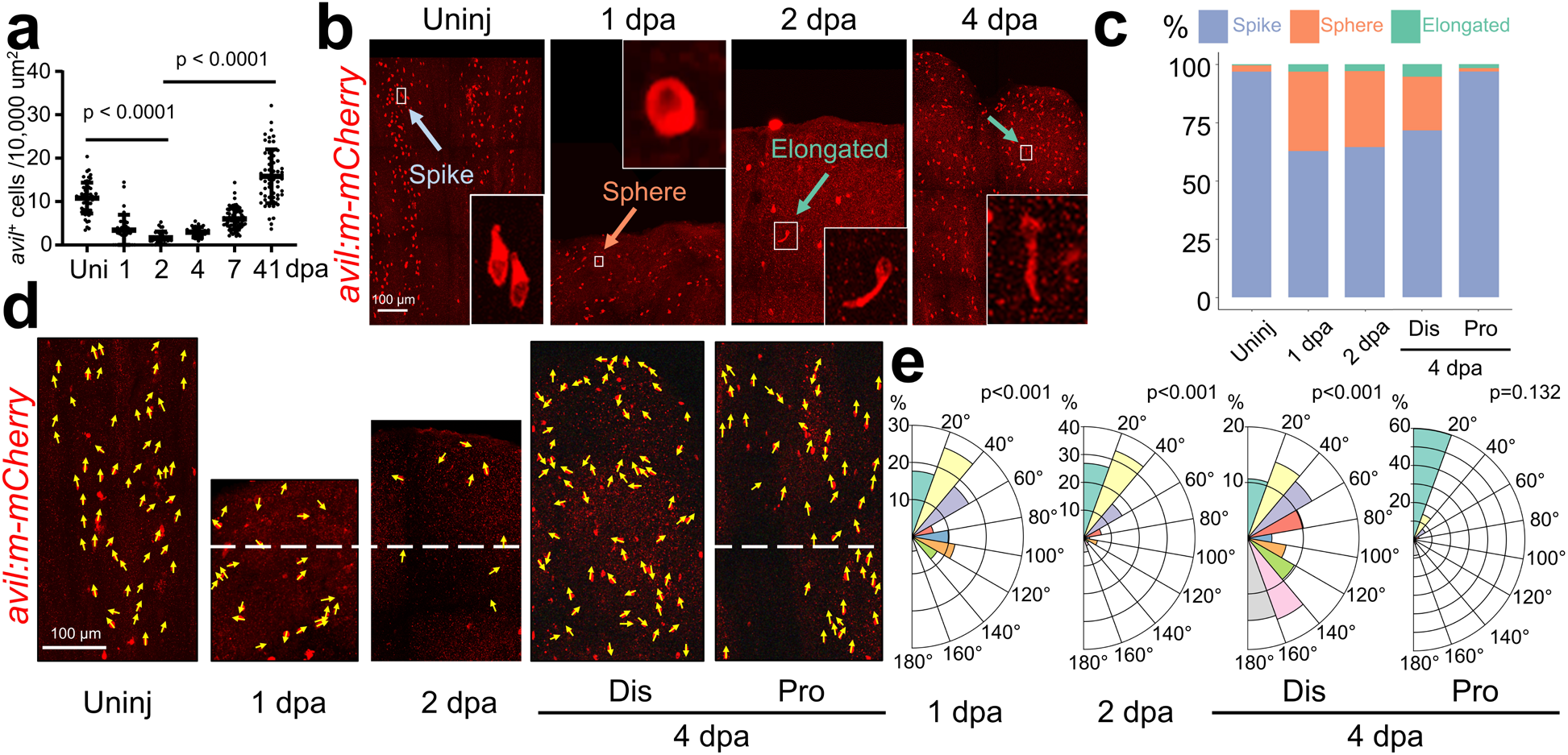
Spike cells exhibit dynamic features during fin regeneration. (**a**) Quantification of *avil*⁺ cell density in uninjured and regenerating fins at 1, 2, 4, 7, and 41 dpa. (n = 56, 36, 36, 36, 66, and 57 regions, respectively, from 6 fish per group). Data are presented with mean ± SD. (**b**) Whole-mount confocal image of *avil:m-mCherry* uninjured and regenerating fins at 1, 2, and 4 dpa. Boxed regions are shown at higher magnification in the insets. (**c**) Quantification of spherical, elongated and spike-shaped *avil*⁺ cells in uninjured and regenerating fins. Total cell counts were 346, 182, 415, 256, and 122, for Uninj, 1 dpa, 2 dpa, 4 dpa-distal (Dis), and 4 dpa-proximal (Pro), respectively (18 fin rays from 6 fish for uninjured and 1 and 2 dpa; 9 fin rays from 3 fish for 4 dpa). (**d**) Representative whole-mount fluorescence images of *avil*⁺ cells in uninjured and regenerating fins at 1, 2, and 4 dpa. Arrows represent the orientation of spike cells. White dashed lines indicate the amputation plane. (**e**) Rose plots depicting the orientation of spike cells, with 0° and 180° representing D and P directions, respectively. Uninjured rose plot is shown in Fig. 2f. Total cell counts were 51, 138, 413, and 256 cells for 1 dpa, 2 dpa, 4 dpa-Dis, and 4 dpa-Pro, respectively (9 fin rays from 3 fish). The proportions of cells oriented within 0–20° were analyzed using Pearson’s chi-square test followed by two-sided Fisher’s exact tests comparing each condition with the uninjured group (Fig. 2f), with Holm correction for multiple comparisons. Adjusted P values are indicated in the panels. Scale bars: **b**, **d**, 100 µm.

Intriguingly, we observed dynamic changes in *avil*^+^ cell morphology during fin regeneration (**Fig. 4b, c**). In uninjured fins, nearly all *avil*^+^ cells displayed their characteristic spike morphology, with only a small minority exhibiting spherical or elongated morphologies. In contrast, regenerating fins showed an elevated number of sphere-shaped and elongated *avil*^+^ cells. Following fin amputation, the proportion of spike-shaped *avil*^+^ cells dropped to 62.6% and 64.3% at 1 and 2 dpa, respectively. By contrast, spherical populations increased markedly to 34.1% and 32.8% at 1 and 2 dpa, respectively. Elongated cells also increased significantly to 3.3% and 2.9% at 1 and 2 dpa, respectively (n = 182 and 415 for 1 and 2 dpa, respectively). During the outgrowth phase of fin regeneration, typically after 2 dpa, distal tip (Dis) regions remain proliferative, whereas the proximal region (Pro) near the amputation plane undergoes redifferentiation^72^. At 4 dpa, the Dis region retained high proportions of spherical and elongated cells similar to 1 and 2 dpa fins (71.5%, 23.1%, and 5.5% for spike, spherical, and elongated cells, respectively; n = 256), whereas the Pro redifferentiation zone more closely resembled uninjured fins (96.7%, 1.6%, and 1.6% for spike, spherical, and elongated cells, respectively; n = 122) (**Fig. 4c**). These results demonstrate dynamic changes in the distribution of *avil*⁺ cell morphologies during regeneration. Spherical and elongated morphologies increase in frequency during early regeneration and in the Dis region of the regenerate, whereas the characteristic spike morphology predominates in the Pro redifferentiation region. These population-level observations suggest that distinct *avil*⁺ cell morphologies are associated with different stages and spatial regions of fin regeneration.

We next investigate whether the P-to-D orientation of *avil*^+^ spike cells is affected during fin regeneration. Analysis of spike cell orientation revealed a population-level reduction in alignment with the P-to-D axis at 1 and 2 dpa. The proportion of spike cells oriented within 0 – 20° of the P-to-D axis was significantly reduced, accompanied by a shift toward orientations greater than 20° (**Fig. 4d, e** and **S3b, c**). At 4 dpa, the spike cell population in the Dis region of regenerate showed weak overall alignment and no strong directional bias (**Fig. 4d, e** and **S3b, c**). In contrast, the Pro redifferentiation region showed a strong population-level bias toward the P-to-D axis (**Fig. 4d, e** and **S3b, c**). Together, these results demonstrate that the population-level distributions of both spike cell morphology and orientation vary across different stages and spatial regions of fin regeneration.

### EdU pulse-chase tracing identifies *de novo* formation and progressive morphological differentiation of *avil*^+^ cells into spike cells

Lost cells after injury can be replenished through distinct mechanisms, including stem cell/progenitor-mediated regeneration and proliferation of pre-existing cells^74^. To distinguish between *de novo* formation and proliferation of pre-existing cells underlying spike cell replenishment, we performed an EdU pulse-chase experiment to label newly divided cells and track their contribution to spike cell regeneration (**Fig. 5a**). A single pulse of the thymidine analog EdU was administered to *avil:l-Green* fish via intraperitoneal (IP) injection at 2 dpa, the time point when *avil^+^* cell numbers begin to increase markedly (**Fig. 4a**). Fins were collected at 2, 12, 36, and 60 hours post-treatment (hpt), and EdU^+^ and *avil:l-Green*^+^ cells were analyzed. At 2 hpt, none of the *avil:l-Green*^+^ cells incorporated EdU (**Fig. 5b, c**), indicating minimal proliferative activity among pre-existing *avil⁺* cells. By 12 and 36 hpt, 2.3% and 5.0% of *avil:l-Green*^+^ cells were EdU^+^, respectively (**Fig. 5b, c**). This proportion further increased to 16.4% by 60 hpt (**Fig. 5b, c**), indicating a temporal delay between EdU labeling and the appearance of EdU^+^ *avil:l-Green*^+^ cells.

**Figure 5.**
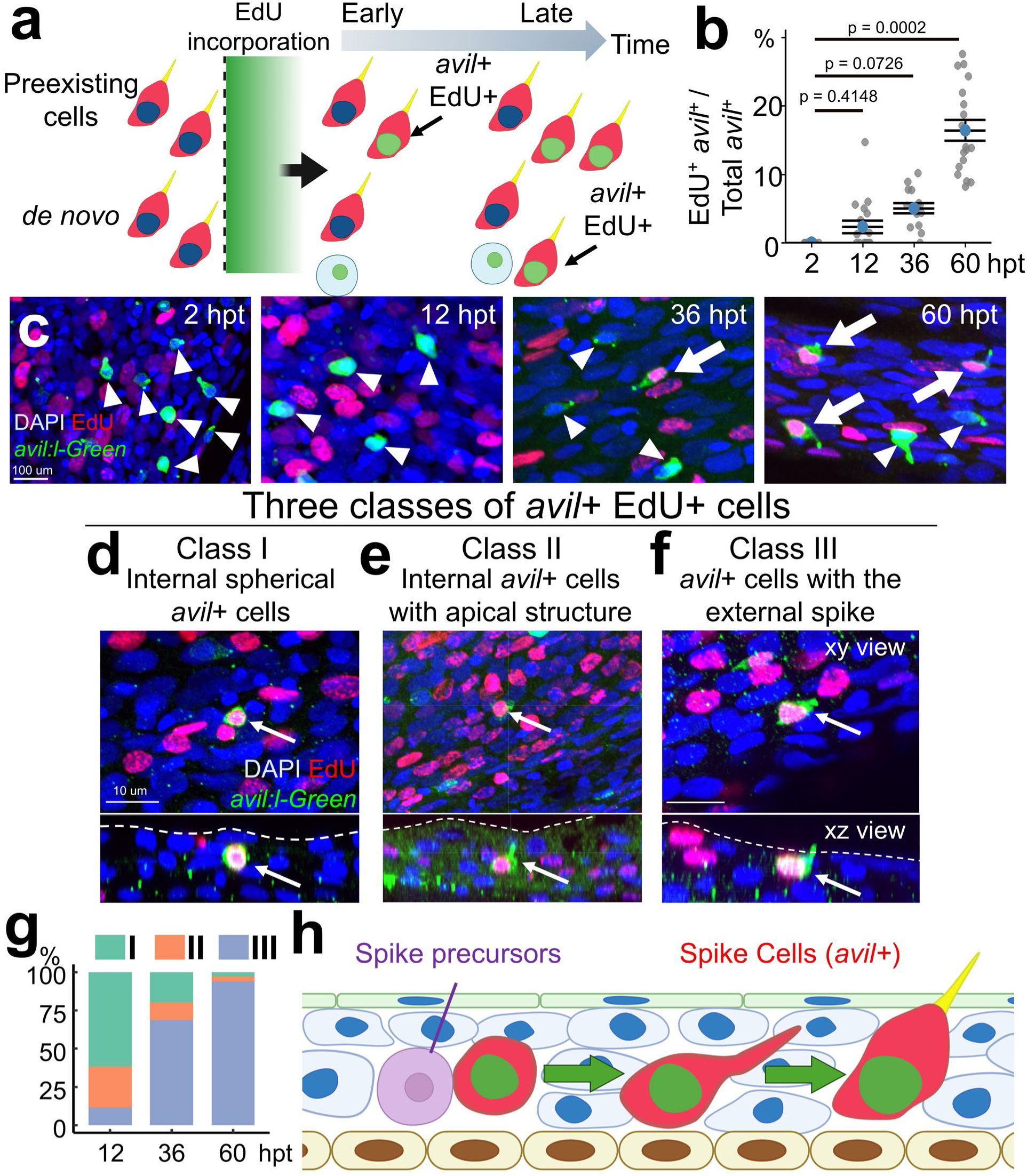
EdU pulse–chase experiments reveal a delayed emergence of EdU-incorporated *avil*⁺ spike cells. (**a**) Schematic illustrating EdU pulse–chase labeling to distinguish self-renewing versus stem/progenitor-derived models of *avil^+^* spike cell generation. (**b**) Following EdU incorporation, *avil:lifeact-mNeongreen (avil:l-Green)^+^* cells were scored as EdU⁺ or EdU⁻ at successive time points. The ratio of EdU⁺ *avil:l-Green^+^* cells to total *avil:l-Green^+^* cells was quantified at 2, 12, 36, and 60 hours post-treatment (hpt) (n = 18, 18, 15, and 18 regions from 6, 6, 5, and 6 fish, respectively). Statistical comparisons were performed using the Kruskal–Wallis test followed by Dunn’s multiple-comparisons test with Holm correction. Adjusted P values are indicated. Data are presented with mean ± SEM. (**c**) Representative confocal images showing EdU incorporation in *avil:l-Green^+^* cells at 2, 12, 36, and 60 hpt. Arrows indicate EdU⁺ *avil:l-Green^+^* cells, whereas arrowheads mark EdU⁻ *avil:l-Green^+^* cells. (**d - f**) Classification of EdU⁺ *avil:l-Green^+^* cells into three morphological categories based on cellular position and structure: internal spherical *avil*⁺ cells, internal *avil*⁺ cells with apical structures, and *avil*⁺ cells bearing an external spike. Representative XY and XZ views are shown. (**g**) Quantification of the relative distribution of the three EdU⁺ *avil:l-Green^+^* cell classes at 12, 36, and 60 hpt (n = 26, 61, and 215 cells, respectively, from 6, 5, and 6 fish). (**h**) Working model illustrating the delayed differentiation of EdU-labeled epithelial stem/progenitor cells into mature *avil*⁺ spike cells during fin regeneration. Scale bars: **c,** 100 µm; **d-f,** 10 µm.

To comprehensively analyze the behavior of newly formed *avil:l-Green*^+^ cells during spike cell morphogenesis, we next focused on EdU^+^ *avil:l-Green*^+^ cells and assessed their morphology and position. We identified three distinct categories of EdU^+^ *avil:l-Green*^+^ cells (**Fig. 5d-g**). Class I consisted of spherical cells located primarily in suprabasal layers corresponding to the 2^nd^ or 3^rd^ intermediate epidermis layers (**Fig. 5d and Supplementary Movie 4**). Class II cells were also located in the suprabasal layers but displayed an apical extension (**Fig. 5e and Supplementary Movie 5**). Class III cells expand their cellular area beneath the superficial layer and form a prominent spike structure that penetrates the superficial epithelium and protrudes externally, representing fully shaped spike cells (**Fig. 5f and Supplementary Movie 6**).

Class I, the internal spherical population, constitutes the predominant category at 12 hpt (61.54%) but progressively declines in proportion at 36 and 60 hpt. Class II cells showed a similar decline. In contrast, the proportion of Class III spike cells increased markedly over time, becoming the dominant population by 60 hpt (**Fig. 5g**). Together, these data indicate that newly formed spike cells arise *de novo* from proliferative precursors located at the basal or suprabasal layer. Their progeny initially lack apical projections and progressively differentiate to acquire the characteristic spike morphology during regeneration (**Fig. 5h**).

### Heterogeneous specialized epithelial cell populations in zebrafish epidermis

We designated the “*avil*-enriched” population because it displayed elevated *avil* expression at the cluster level. At the single-cell level, however, *avil* transcripts were detected in only 19.8% of cells in the cluster. It raised questions about the cellular identity and heterogeneity of the *avil*-enriched population. To address these questions, we performed subclustering analysis of this population in the M-fins scRNA dataset (**Fig. 1e, asterisk**) and identified 9 transcriptionally distinct subclusters (**Fig. 6a and S5a**). *pou2f3*, a transcription factor associated with tuft cells in mammals^62^, was broadly expressed across most clusters except Cluster 6 (**Fig. S5b**). In contrast, *avil* was most strongly enriched in Cluster 2, with modest expression in subsets of cells in Clusters 0, 1, 4, and 7, but no detectable expression in Clusters 3, 5, and 6 (**Fig. 6c**). Based on our findings that *avil* marks mature spike cells, Cluster 2 likely represents differentiated spike cells (**Fig. 2, 3, 4, 5, 6b**). Consistently, *gng13a*, a homolog of mammalian *Gng13,* which is a critical signaling component in mammalian Tuft cells^79^, was also enriched in these clusters, including Cluster 2, along with *crp7, rgs4,* and *spry2* (**Fig. S5c**).

**Figure 6.**
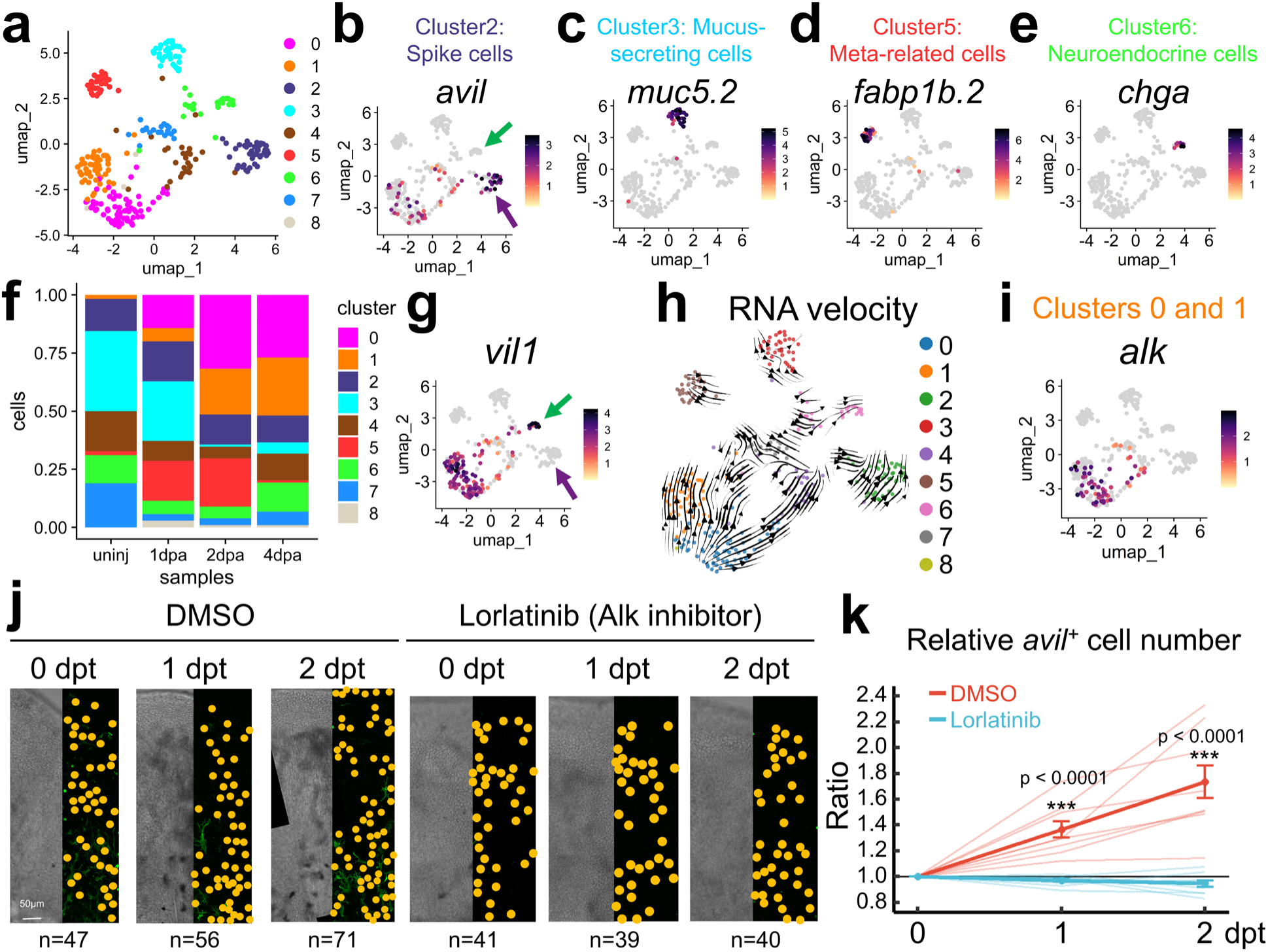
Epithelial subclustering identifies Alk-dependent spike cell replenishment from stem/progenitor cells during fin regeneration. (**a**) UMAP plot of *avil*-enriched Epi subclusters from the M-fins dataset. (**b**) *avil*, a marker for spike cells, shows strongest expression in Cluster 2. (**c**) Expression of the mucin gene *muc5.2* in Cluster 3, indicating goblet cells. (**d**) *fabp1b.2*, a fatty acid binding protein, is enriched in Cluster 5, indicating metabolism (Meta)-related cells. (**e**) Neuroendocrine genes (*chga*) are highly enriched in Cluster 6. (**f**) Proportion of 9 subclusters across different time points. (**g**) UMAP feature plot of *vil1* expression. (**h**) RNA velocity stream plot. (**i**) UMAP feature plot of *alk* expression. (**j**) Representative whole-mount confocal images showing the distribution of *avil*⁺ spike cells in regenerating fins treated with DMSO or Lorlatinib at 0, 1, and 2 dpt. *avil^+^* cells are marked by yellow dots. Full images of regenerating fins are shown in **Fig. S7b**. (**k**) Quantification of spike cell numbers over time, shown as the ratio of total *avil*⁺ cell numbers at 1 and 2 dpt relative to 0 dpt for individual fins (n = 9 and 10 fish for DMSO and Lorlatinib, respectively). Lorlatinib treatment markedly attenuates spike cell replenishment compared with DMSO controls. Statistical significance was determined using a linear mixed-effects model with treatment, time, and their interaction as fixed effects and fish identity as a random effect, followed by pairwise comparisons with Holm correction. Data are presented with mean ± SEM. Scale bars: **j,** 50 µm.

Other subclusters exhibited transcriptional signatures consistent with distinct specialized epithelial cell populations. Cluster 3 was highly enriched for *mucin 5.1 (muc5.1), muc5.2,* and *creb3l1/oasis*, consistent with a mucus-secreting goblet cell identity (**Fig. 6b and S5c**)^14, 80, 81^. Cluster 5 was characterized by the expression of metabolism-related genes, including *fabp1b.2, slc16a12b, and gstm.1*, (**Fig. 6c and S5c**). Cluster 6 was labelled by *chga, cartpt, scgn, scg3,* and *penkb*, genes encoding hormones or secreted peptides, suggesting a neuroendocrine- or enteroendocrine-like identity (**Fig. 6d and S5c**)^82, 83^. Together, these analyses indicate that the initially defined *avil*-enriched population is transcriptionally heterogeneous and contains multiple specialized epithelial populations.

We also observed dynamic changes in subcluster composition across time points (**Fig. 6f and S5a**). Clusters 3 and 7 were abundant in uninjured tissues but declined during fin regeneration. Clusters 2, 4, and 6 were present across all time points. Cluster 5 transiently expands at 1 and 2 dpa but returns to the uninjured level by 4 dpa. Interestingly, Clusters 0 and 1, two closely related populations, emerged robustly following fin amputation (**Fig. 6f and S5a**), suggesting potential injury-responsive roles. *avil* and *villin* belong to the Gelsolin/Villin superfamily of actin-binding proteins but mark distinct intestinal epithelial cell types in mammals: *Avil* specifically labels tuft cells, while *Villin* is expressed in other differentiated epithelial lineages^84, 85^. Consistent with this, *villin1* (*vil1*), the zebrafish ortholog of *Villin*, and *avil* distinctly marked differentiated populations in Clusters 6 and 2, respectively (Arrows in **Fig. 6b and 6g**). However, cells in Clusters 0 and 1 exhibit moderate expression of both genes. Given their injury-associated emergence, Clusters 0 and 1 may represent stem/progenitor populations that give rise to Clusters 2 and 6. To investigate the lineage dynamics of these specialized epithelial cells, we performed RNA velocity analysis, which revealed directional flow from Clusters 0 and 1 toward Clusters 2 and 6 (**Fig. 6h**). These results suggest that Clusters 0 and 1 represent precursor populations that contribute to spike cell generation during fin regeneration.

We next examined genes highly enriched in Clusters 0 and 1 and identified *anaplastic lymphoma kinase (alk)*, a receptor tyrosine kinase associated with cell proliferation and stemness ^86^, along with *aldh1a3, rbp5, stra6,* and *ehf* as distinctive markers of these clusters (**Fig. 6i and S5c**). To determine whether spike cells and the *alk*-enriched epithelial population are identified in an independent dataset, we reanalyzed a single-nucleus RNA-seq dataset of postembryonic zebrafish trunk skin ^87^ (**Fig. S6**). Subclustering of specialized epithelial populations, including ionocytes, goblet cells, and MLCs, revealed a cluster enriched for *avil* and *spry2*, consistent with the transcriptional identity of spike cells (**Fig. S6a–c**). Notably, we also identified a distinct epithelial population enriched for *alk* and *aldh1a3*, resembling the *alk*-enriched population present in M-fins scRNA dataset (**Fig. S6b, d**). These findings provide independent transcriptomic support for the presence of both spike cells and *alk*-enriched epithelial populations in zebrafish trunk skin.

To investigate the functional role of Alk signaling in spike cell replenishment, we treated *avil:l-Green* zebrafish with vehicle control and Lorlatinib, a small-molecule ALK/ROS1 tyrosine kinase inhibitor^88^. In zebrafish, pharmacological inhibition with Lorlatinib causes iridophore defects that phenocopy loss of *ltk* and *alkal* ligand genes, indicating suppression of Alk-family signaling ^89^. Treatment was administered from 3 to 5 dpa, and the total number of *avil:l-Green*^+^ cells was quantified daily (**Fig. S7a**). *avil:l-Green*^+^ cell numbers increased in vehicle-treated control group, consistent with active regeneration. In contrast, *avil:l-Green*^+^ cell numbers within fins of Lorlatinib administered fish were constant without increasing, indicating a substantial reduction in the generation of new *avil:l-Green*^+^ cells (**Fig. 6j, k and S7b**). Lorlatinib treatment was unlikely to affect overall fin regeneration (**Fig. S7c**), indicating that Alk inhibition specifically impairs replenishment of proliferative *avil*^+^ cell populations..

### *avil*⁺ spike cells support the local maintenance of *mpeg1*⁺ immune cells

During development, tissue-resident immune cell populations expand to meet local tissue demands and are subsequently maintained within their tissue niches. This homeostasis is regulated by the local microenvironment through interactions with neighboring cells^90, 91^. Given the close spatial association between *avil*⁺ spike cells and *mpeg1*⁺ immune cells (**Fig. 3**), we questioned whether spike cells contribute to maintenance of the local immune cell pool in the fin epidermis. To this end, we employed the Mtz-NTR genetic ablation system^92^ and generated *avil:BFP-NTR* transgenic fish to specifically deplete *avil*^+^ spike cells. Treatment with 100 µM Mtz for 12 hours efficiently ablated *avil:BFP-NTR^+^* spike cells: *avil:BFP-NTR^+^* cell numbers decreased sharply by 12 hpt, were nearly eliminated by 36 hpt, and subsequently recovered (**Fig. S8a, b**). These data demonstrate efficient and reversible genetic ablation of spike cells, establishing a platform to assess *mpeg1^+^* cells dynamics in the absence of spike cells.

Using *avil:BFP-NTR;mpeg1:EGFP* transgenic fish, we next quantified *mpeg1^+^* cell abundance following spike cell ablation (**Fig. 7a**). Upon Mtz treatment, *avil:BFP*^+^ cells were robustly depleted at 12 and 36 hpt, respectively. Notably, *mpeg1^+^* cell numbers also declined after spike cell ablation and reached their lowest level at 36 hpt, when spike cell loss was maximal. Specifically, *mpeg1⁺* cell abundance decreased to 77.1% and 62.5% of pre-ablation levels at 12 and 36 hpt, respectively (**Fig. 7a-d**). As *avil:BFP*⁺ spike cells gradually recovered, *mpeg1*⁺ cell numbers also increased toward pre-ablation levels (**Fig. 7c, d and S8d, e**). To visualize the dynamic association between spike cells and *mpeg1*⁺ cells during recovery, we performed longitudinal whole-mount confocal imaging of *avil:m-mCherry; avil:BFP-NTRv2; mpeg1:EGFP* triple-transgenic fish. Consistent with our quantitative analyses, spike cell depletion led to a reduction in *mpeg1^+^* cell density. As newly regenerated *avil*⁺ spike cells emerged, close contacts between spike and *mpeg1*⁺ cells were observed as early as 60 hpt (**Fig. 7e**). Spatial analysis showed that *avil*⁺ cells were enriched in intra-fin ray relative to inter-fin ray regions, whereas *mpeg1*⁺ cells were distributed across both regions, with modestly higher abundance in inter-fin ray regions (**Fig. S8c**). These regional distribution patterns were re-established following recovery from spike cell ablation. These findings indicate that maintenance of the immune niche within the fin epidermis is closely associated with the presence of *avil*⁺ spike cells.

**Figure 7.**
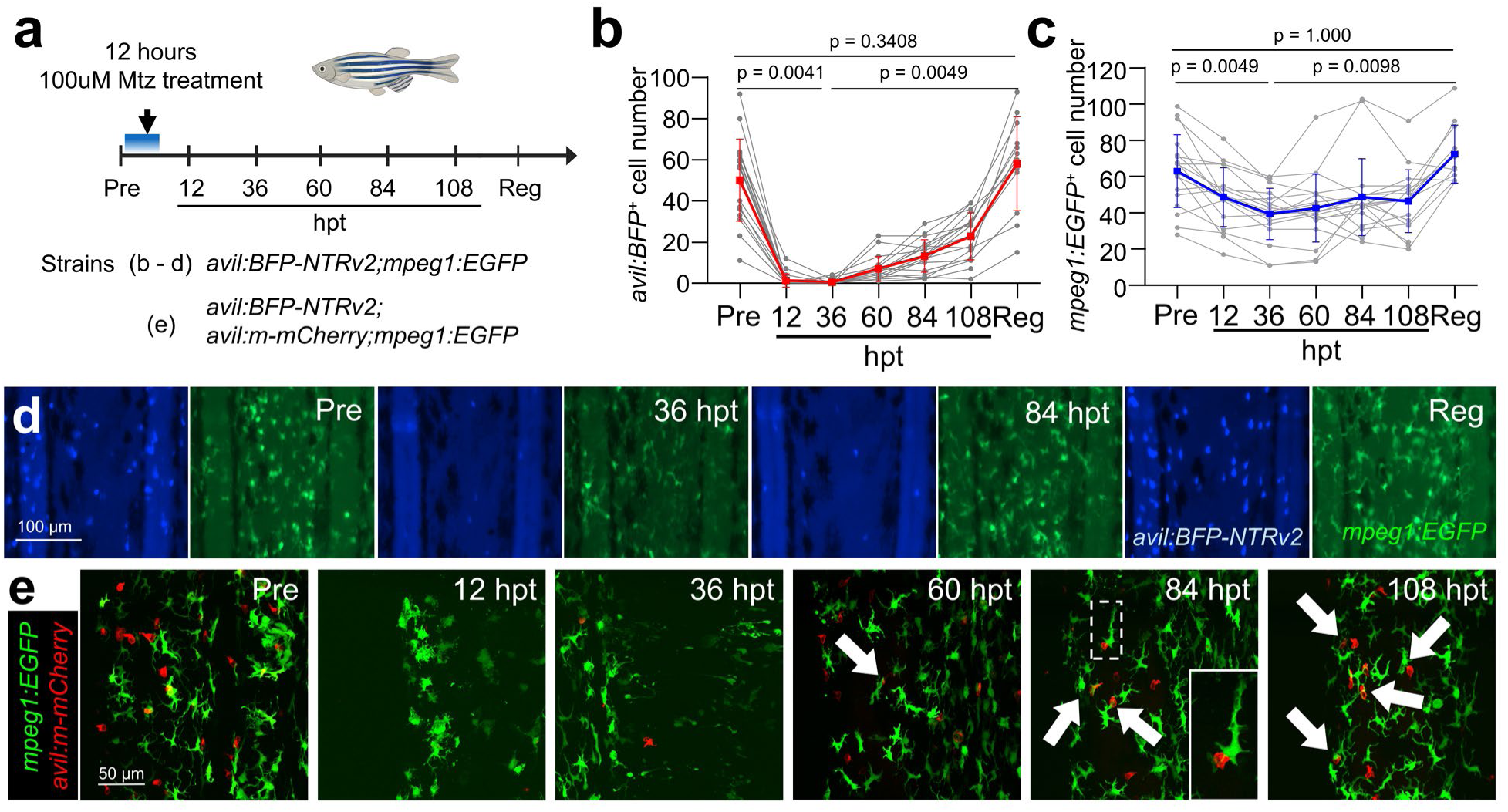
*avil*⁺ spike cells regulate the resident *mpeg1*^+^ immune cells pool. (**a**) Experimental scheme illustrating 12-hour Mtz treatment in *avil:BFP-NTRv2;mpeg1:EGFP* fish, followed by fluorescence imaging analysis before treatment (Pre) and at 12, 36, 60, 84, and 108 hpt, and at the recovery >14 dpt (Reg). Strains used for the respective panels are indicated. (**b**) Quantification of relative *avil:BFP⁺* cell number in *avil:BFP-NTRv2;mpeg1:EGFP* zebrafish, normalized to the pre-treatment level. (**c**) Quantification of relative *mpeg1:EGFP⁺* immune cell number in *avil:BFP-NTRv2;mpeg1:EGFP* zebrafish, normalized to the pre-treatment level. Data was obtained from 18 fin rays from 6 fish for Pre and 12, 36, 60, 84, and 108 hpt and 12 fin rays from 4 fish for Reg. Statistical significance was assessed using Friedman repeated-measures tests, followed by paired Wilcoxon signed-rank tests with Holm correction for planned comparisons. Data are presented with mean ± SEM. (**d**) Representative fluorescence images of *avil:BFP-NTRv2;mpeg1:EGFP* fins at Pre, 36 and 84 hpt, and Reg. (**e**) Representative confocal images of *avil:BFP-NTRv2;avil:m-mCherry;mpeg1:EGFP* fins at the indicated time point. White arrowheads indicate contact regions between recovered *avil:m-mCherry⁺* cells and *mpeg1:EGFP⁺* immune cells. The boxed region at 84 hpt is shown at higher magnification in the inset. Scale bars: **d,** 100 µm; **e,** 50 µm.

### Spike cells maintain a local immune niche with dynamic cell association

Immune cells play essential roles in intercellular communication, which is often facilitated by their migratory behavior ^74, 93, 94^. Previous studies showed that *mpeg1*^+^ cells in zebrafish skin are motile ^49, 75, 76^, suggesting that their interaction with spike cells may be dynamic. To investigate the dynamics and persistence of these interactions, we developed a photoconversion-based cell-tracking strategy (**Fig. 8a**). We crossed photoconvertible *mpeg1:Dendra2* transgenic fish ^95^ with the *avil:BFP-NTR* line to obtain *avil:BFP-NTR; mpeg1:Dendra2,* enabling individual *mpeg1*⁺ cells to be labeled and tracked over an extended period. We selectively photoconverted a small number of *mpeg1*⁺ cells from green to red (*mpeg1:G->R*) within a single fin-ray bone segment (**Fig. 8b**) and subsequently performed daily longitudinal confocal imaging (**Fig. 8c**). This approach allowed us to track individual photoconverted *mpeg1:G->R* cells and monitor their spatial relationships with *avil*⁺ spike cells over time.

**Figure 8.**
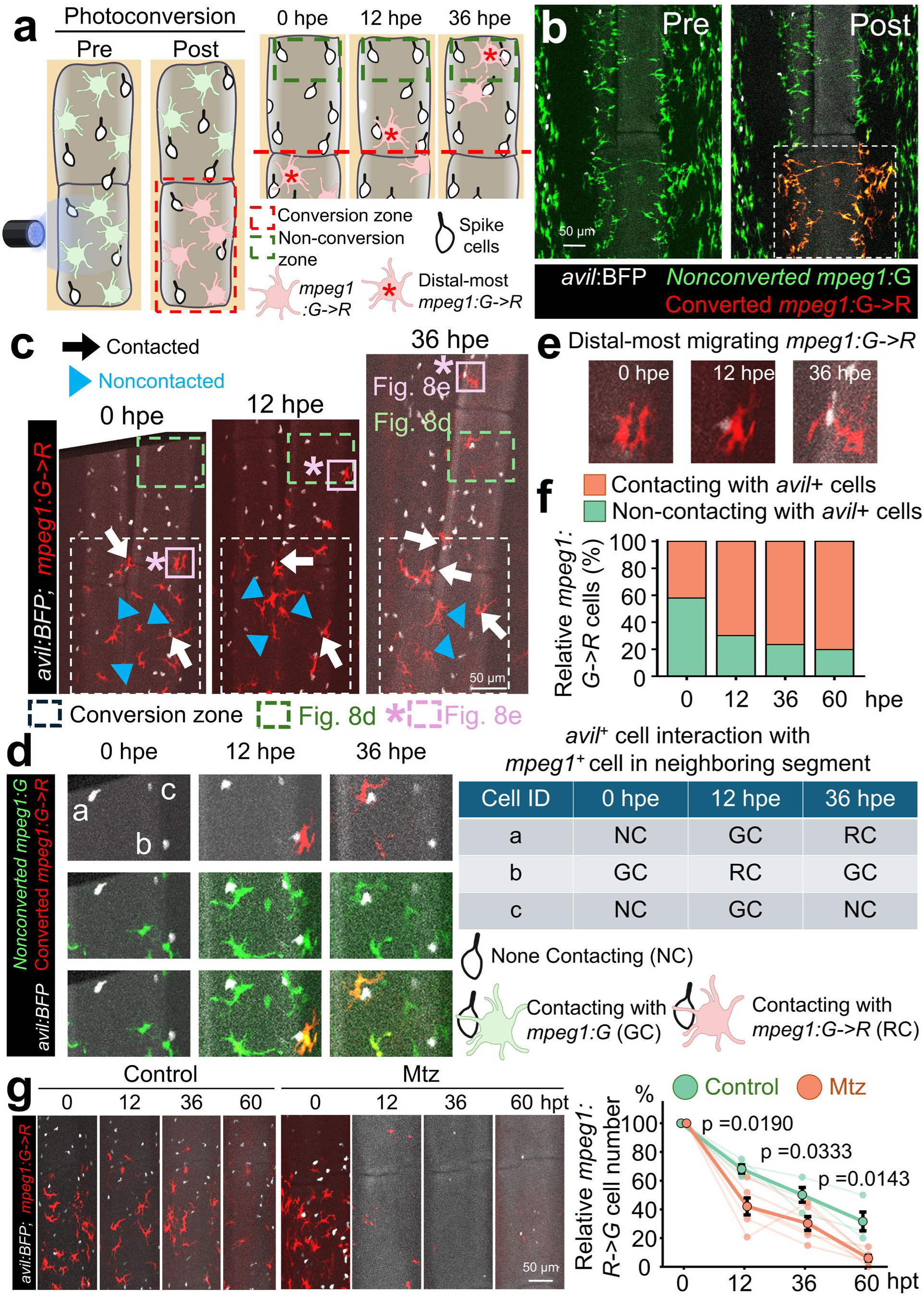
Photoconversion-based tracking reveals dynamic interactions between *avil*⁺ spike cells and *mpeg1*⁺ immune cells. (**a**) Schematic of the photoconversion-based cell tracking strategy. Photoconversion switches Dendra2 fluorescence from green to red in *mpeg1^+^* cells, *mpeg1:G->R,* within a single intra-fin ray bone segment. Confocal imaging captured the photoconverted segment (red dashed box, conversion zone) and the adjacent distal segment (green dashed box, non-conversion zone). hpe, hours post-exposure. (**b**) Representative confocal images of *avil:BFP;mpeg1:Dendra2* fins before (Pre) and after (Post) photoconversion. The dashed box indicates the photoconverted region. (**c**) Representative images displaying *avil^+^* spike cells (white) and photoconverted *mpeg1:G->R* cells at 0, 12, and 36 hpe. White arrows indicate *mpeg1:G->R* contacting *avil*⁺ spike cells, whereas blue arrowheads indicate non-contacting *mpeg1:G->R*. White dashed boxes indicate the photoconversion zone. Green and pink dashed boxes are regions shown in **d** and **e**, respectively. (**d**) Representative images showing changes in the contact status of individual *avil*⁺ spike cells (a–c) with *mpeg1*⁺ immune cells in the adjacent non-conversion zone. The table summarizes their contact status at 0, 12, and 36 hpe. NC, non-contacting; GC, contacting with non-photoconverted *mpeg1:G* cells; and RC, contacting with photoconverted *mpeg1:G->R* cells. (**e**) Representative images of a photoconverted *mpeg1:G->R* cell tracked near the distal edge at 0, 12, and 36 hpe. (**f**) Quantification of relative numbers of *mpeg1:G->R* cells contacting or non-contacting with *avil*⁺ spike cells at 0, 12, 36, and 60 hpe (n = 4 fish). (**g**) Representative images exhibiting *mpeg1:G->R* cells remaining in control and Mtz-treated fish at 0, 12, 36, and 60 hpt. Cell counts were normalized to the corresponding baseline count (0 h) for each fish (n = 4 and 6 fish for control and Mtz, respectively). Statistical significance was assessed using two-sided exact permutation tests with Holm correction for multiple comparisons. Data are presented with mean ± SEM. Scale bars: **b**, **c**, **g**, 50 µm.

The adjacent intra-fin ray bone segment, designated the non-conversion zone, contained no *mpeg1:G->R* at 0 hour post-exposure (hpe). Longitudinal imaging revealed that some *mpeg1:G->R* cells migrated from the conversion zone into the adjacent non-conversion zone at 12 and 36 hpe (**Fig. 8c**). In contrast, *avil*^+^ cells remained in the same location, suggesting that spike cells are unlikely to migrate under homeostatic conditions.

We first examined individual *avil*⁺ spike cells in the non-conversion zone to assess their contacts with *mpeg1*^+^ cells. Interestingly, these observations revealed dynamic changes in contacting partners, including transitions between non-contacting states (NC) and contacts with either non-photoconverted *mpeg1:G* (GC) or photoconverted *mpeg1:G->R* (RC) cells (**Fig. 8d and S9**). We next focused on *mpeg1:G->R* cells that migrated to the most distal region (**Fig. 8c, e**). These distally located *mpeg1:G->R* cells were observed not only farther from the conversion zone but also in close proximity to *avil⁺* spike cells, where they established contacts. We also quantified the proportion of photoconverted *mpeg1:G->R* cells contacting *avil*⁺ spike cells during longitudinal imaging. The contacting fraction progressively increased from 42.0% at 0 hpe to 80.2% at 60 hpe, accompanied by a corresponding decrease in the non-contacting fraction (**Fig. 8f**). Collectively, these observations indicate that association between spike and immune cells is highly dynamic within the fin epidermis rather than remaining fixed between individual cell pairs.

To determine whether spike cells contribute to the local persistence of *mpeg1^+^* immune cells, we combined photoconversion-based tracking with Mtz-mediated *avil*⁺ cell ablation. We followed the photoconverted *mpeg1:G->R* cell population in control and Mtz-treated fish and quantified the proportion of cells (**Fig. 8g**). Compared with controls, Mtz-treated fish exhibited a substantial decline in the number of remaining *mpeg1:G->R* cells at 12, 36, and 60 hpe (**Fig. 8g**). These findings suggest that *avil*⁺ spike cells contribute to the local retention of *mpeg1*⁺ immune cells. Overall, our results suggest that spike cells support the maintenance of a local immune cell population while immune cells remain motile and dynamically remodel their contacts.

## Discussion

Here, we define a specialized epithelial cell type in the zebrafish fin epidermis. *avil*-expressing cells exhibit a distinctive morphology characterized by an apical spike structure that protrudes outward. Their molecular signature, enriched for *avil* and *pou2f3*^22, 24, 62^, suggests that these *avil*^+^ cells are analogous to mammalian tuft cells, which are named for their apical tuft-like projections of long, rigid microvilli^21^. Accordingly, we refer to these *avil*⁺ cells as "spike cells" based on their unique apical morphology.

We propose that spike cell maturation proceeds through a sequential trajectory. Upon injury, precursor cells, presumably epithelial stem/progenitor cells, are activated to generate new cells. A subset of these newly formed cells becomes committed to the spike cell fate. Our scRNA-seq and pharmacological analyses indicate that Alk signaling pathway is involved in these early processes. These spike cell-committed cells initially display a spherical morphology lacking an apical projection and are positioned within the suprabasal layer. During subsequent maturation, they increase in size and extend from deeper suprabasal regions toward more superficial epithelial layers. They then develop elongated, actin-rich apical projections that extend outward from the epithelial surface.

Ultimately, they acquire the characteristic morphology of mature spike cells. Similar morphological change during maturation is also observed in intestinal organoid model^96^, suggesting evolutionary conserved features of tuft/spike cell maturation.

Solitary chemosensory cells (SCCs) are individual sensory cells broadly distributed in aquatic vertebrates, including teleost fish^97^. SCCs reside in various epithelial tissues, such as the skin, gill, and oropharyngeal cavity, and presumably function as environmental sensors^98, 99^. Given that SCCs typically exhibit a single apical spike-like projection^100^, the *avil*^+^ spike cells analyzed in this study likely represent SCCs. While ultrastructural features of SCCs have been described using transmission electron microscopy^100, 101^, their cellular characteristics visualized by fluorescence microscopy remain largely unexplored. Here, we generate transgenic reporter lines to analyze spike cells, potentially corresponding to SCCs, and uncovered notable features. First, spike cells are large, spanning nearly the full thickness of the entire epidermis from the surface to the suprabasal layer. Second, their apical structures protrude beyond the surface squamous epithelial cells with consistent P-to-D orientation in caudal fins. These protrusions emerge at various junctions between squamous epithelial cells, including both side boundaries and vertex points. Third, the spike protrusions are highly enriched in F-actin, indicating that spike cells possess distinct cytoskeletal components. This is reminiscent of mammalian tuft cells, which contain specialized actin-core bundles within their tufts^102^. Together, these findings provide a comprehensive characterization of *avil*⁺ spike cells in the zebrafish fin epidermis and support the idea that they represent an evolutionarily conserved epithelial cell type analogous to mammalian tuft cells.

Studies in mice implicated that epithelial stem cells, such as Lgr5^+^ intestinal stem cells, primarily contribute to tuft cell generation under normal homeostatic conditions ^25, 103, 104^. More recently, human intestinal organoid studies revealed that mature tuft cells can become proliferative in response to specific immunological cues. Notably, a single *AVIL^+^* tuft cell can generate complete organoids comprising all major intestinal epithelial lineages ^24^. Wound-repair-associated cytokines, such as IL-4 and IL-13, can induce dedifferentiation of mature tuft cells into stem cells. Moreover, recent work found that metaplastic tuft cells in pancreatic cancers can transdifferentiate into neural-like progenitor cells^99^, highlighting the context-dependent plasticity of tuft cells. To explore the regenerative mechanisms of zebrafish spike cells, we employed tailfin amputation and examined spike cell dynamics during fin regeneration. Following fin amputation, spike cell numbers initially declined, but recovered robustly beginning at 2 dpa. Our data demonstrates a limited contribution of mature spike cells to the generation of new spike cells. Instead, spike precursors are activated by injury to generate new spike cells *de novo*, indicating a stem-cell/progenitor-mediated mechanism as the primary mode for spike cell replenishment. It would be of interest to explore the conditions under which spike cells can dedifferentiate and investigate their cellular and molecular features influencing this plasticity.

Mammalian tuft cells function as specialized chemosensory epithelial cells that monitor environmental and pathogen-derived cues and mediate immune responses ^5, 19, 20, 22, 23, 27, 28, 29, 31, 32, 33^. These cells act as local sentinels at mucosal surfaces. Beyond these roles, our study suggests that specialized epithelial cells are also important to the maintenance of local immune cell populations. We found that *avil*⁺ spike cells are closely associated with a subset of *mpeg1*^+^ through direct contacts. Notably, due to migratory features of immune cells, association between *avil*^+^ spike cells and *mpeg1*^+^ immune cells is highly dynamic. Loss of spike cells resulted in a significant reduction in *mpeg1*^+^ immune cell abundance, and spike cell recovery contributes to repopulating these cells. Together, these findings support a model in which spike cells serve as niche-forming cells that help sustain resident *mpeg1*^+^ immune cell pools. This phenomenon is conceptually analogous to quorum sensing in bacterial systems, wherein cell-cell communication regulates population dynamics^105^. Further investigation of spike cells using advanced imaging and genetic tools in zebrafish will provide important insights into how specialized epithelial cells coordinate sensory functions with immune regulation.

Overall, our study establishes the zebrafish fin spike cell system as a tractable *in vivo* platform for investigating regeneration of tuft cell-like epithelial population and epithelial–immune interactions. By defining the molecular identity, regenerative origin, maturation process, and *mpeg1*^+^ immune cell-associated behavior of spike cells, we provide a framework for dissecting how specialized epithelial cells are rebuilt after injury and how they help shape local immune responses. Given the optical accessibility and genetic versatility of zebrafish, this system offers a valuable opportunity to study how tuft cell-like epithelial cells coordinate regeneration, immune cell residence, and tissue homeostasis.

## Material And Methods

### Zebrafish

Wild-type or transgenic male and female zebrafish of the outbred Ekkwill (EK) or AB strain ranging up to 18 months of age were used for all zebrafish experiments. The following transgenic lines were used in this study: *Tg(avil:memb-mCherry)^uwk23^*, Tg(avil:lifeact-mNeonGreen)*^uwk51^*, *Tg(avil:BFP-NTRv2)^uwk52^*, *Tg(mpeg1:EGFP)^43^*, *Tg(mpeg1:Dendra2)^uwm12^* ^95^. The water temperature for adult animals was maintained at 26°C. Embryos and larvae were maintained at 28°C in egg water containing 300 mg/L sea salt, 75 mg/L calcium sulfate, 37.5 mg/L sodium bicarbonate, and 0.0001% methylene blue. Animals were anesthetized in 0.02% tricaine until gill movement stopped. Work with zebrafish was performed in accordance with University of Wisconsin-Madison guidelines.

### Subcloning and transgenesis

*avil:memb-mCherry*: pMC-*memb-mCherry-*pA construct was generated by inserting the N-terminal membrane-targeting sequence of zebrafish *gap43* (Gap43-mem) into the pMC-*mCherry*-pA vector via NheI and HindIII enzymes-mediated subcloning. Zebrafish Gap43-mem fluorescent reporters exhibit both plasma membrane and intracellular fluorescence ^66, 67^. A 443-bp genomic sequence immediately upstream of the *avil* start codon and a 512-bp downstream genomic sequence located within the adjacent intron were amplified from zebrafish genomic DNA and sequentially inserted into the *pMC-memb-mCherry-pA* vector using restriction enzyme-based subcloning. The 5’ end of upstream sequence contains modified sgRNA target site (gGGAGATACAGCAGATTTAGAGG). The final minicircle DNA was prepared following the protocol described in ^106^. The corresponding sgRNA was synthesized using a cloning-free method^107^ using the MEGAshortscript T7 Transcription Kit (Invitrogen, AM1354) and purified by the RNA purification Kit (Zymogen, R1016) according to the manufacturer’s instructions. A sgRNA (25∼30 ng/ul) and *avil:memb-mCherry* minicircle (20-25 ng/ul) were mixed and co-injected with Cas9 protein (0.5ug/ul; PNABio, CP01) into the one-cell stage embryos. Larvae expressing mCherry were selected between 3 - 5 days dpf and raised to adulthood, and founders were screened with F_1_ progenies. A total of five independent strains were established, all exhibiting similar expression patterns in the larval olfactory system. We selected lines that exhibited robust spike-cell expression in adult zebrafish fins for further analysis. Genotyping analysis revealed that the injected fragment was randomly integrated rather than precisely targeted to the *avil* locus.

*avil:lifeact-mNeonGreen* and *avil:BFP-NTRv2*: *lifeact-mNeonGreen* was a gift from Dorus Gadella (Addgene #98877 ; http://n2t.net/addgene:98877 ; RRID:Addgene_ 98877)^108^. *mCherry-NTRv2* was a gift from Jeff Mumm (Addgene plasmid # 158653 ; http://n2t.net/addgene:158653 ; RRID:Addgene_158653)^92^. *lifeact-mNeonGreen* was subcloned into pMC-ME vector to generate *pMC-ME-lifeact-mNeonGreen.* For the *BFP-NTRv2* construct, the *mCherry-NTRv2* cassette was first subcloned into a pMC-ME vector, followed by replacement of *mCherry* with *BFP* using overlapping PCR and Gibson assembly, generating *pMC-ME-BFP-NTRv2*. A 443-bp upstream and a 512-bp downstream genomic fragment of the *avil* locus were amplified from the *avil:memb-mCherry* plasmid, each incorporating BsaI recognition and cleavage sites for Golden Gate assembly. To generate the final construct, the upstream and downstream homology arms were ligated with the *pMC-ME-lifeact-mNeonGreen* or *pMC-ME-BFP-NTRv2* cassette into the destination vector pDest-BsaI-Tol2-I-SceI (Addgene# 241214)^109^ using a Golden Gate assembly reaction. The reaction mixture (10 µl total volume) contained 1 µl of T4 DNA ligase buffer (NEB), 25–50 ng of each homology arm cassette, 100 ng of the middle entry vector (*pMC-ME-lifeact-mNeonGreen or pMC-ME-BFP-NTRv2*), 50–100 ng of the destination vector, 0.5 µl of BsaI-HFv2, and 0.5 µl of T4 DNA ligase, with nuclease-free water added to adjust the final volume. The *avil:lifeact-mNeonGreen* or *avil:BFP-NTRv2* construct (25–30 ng/μl) was used as the DNA template and co-injected with I-SceI meganuclease (NEB, R0694L) into one-cell stage embryos to generate transgenic lines via random genomic integration. Larvae expressing fluorescent signals were screened between 3–5 days dpf and raised to adulthood. Founders were identified by screening F_1_ progenies for larval olfactory expression. Transgene-positive F_1_ animals were raised and outcrossed with wild-type to generate F_2_ stable lines. Stable expression of the *avil* reporter was confirmed in F_2_ animals by examining reporter-positive cells in adult caudal fins, and lines showing the expected *avil* expression pattern in the adult fin epidermis were used for subsequent experiments. The sequences of the primers used for subcloning are listed in **Table S1**.

### Whole-mount immunostaining, histology, and imaging

For whole-mount fin immunostaining, fins were fixed with 4% paraformaldehyde (PFA) overnight at 4 °C. After fixation, fins were washed several times in 0.3% Triton X-100 in phosphate-buffered saline (PBST0.3) at room temperature. After that, fins were transferred to blocking solution (1% BSA and 0.3% Triton X-100) for 1 h at room temperature and treated with primary antibody overnight at 4 °C. The next day, fins were washed and incubated with secondary antibodies at 4 °C overnight. To visualize actin filament, fins were stained with fluorescently labeled phalloidin conjugates, including Alexa Fluor™ Plus 405 Phalloidin (Invitrogen, A30104) and Phalloidin-iFluor 488 Conjugate (Cayman Chemical, 20549). The primary and secondary antibodies used in this study were: anti-Ds-Red (rabbit, 632496, Takara Bio, 1:500), Alexa Fluor 594 (mouse and rabbit, A11032 and A11037, Life Technologies; 1:500). Tissues were washed and embedded in mounting medium (Fluoromount-G, 00-4958-02, Thermo Fisher). Whole-mount fin images were acquired using an Olympus Inverted FV3000 Confocal Laser Scanning Microscope (Evident) or an AxioZoom stereo fluorescence microscope (Zeiss) at the time points indicated. Image stitching was automatically processed using Olympus software during image acquisition. Maximum-intensity projections were generated from 60–90 optical sections acquired at 1-µm intervals. For live and longitudinal confocal imaging experiments, individual Z-stacks were registered using StackReg in Fiji/ImageJ with a translation transformation to correct minor image drift. Registration was performed separately for each imaging time point, followed by maximum-intensity projection for visualization. Further image processing was carried out using Adobe Photoshop, FIJI/ImageJ or Imaris (Oxford Instruments) image analysis software.

### Scanning electron microscopy (SEM)

For scanning electron microscopy, zebrafish caudal fins were fixed in 4% PFA overnight at 4 °C. After fixation, samples were washed several times in PBST 0.3 at room temperature. Specimens were then dehydrated through a graded ethanol series and dried by critical point drying using liquid CO₂. Dried samples were mounted on aluminum stubs and sputter-coated with gold (or Au/Pd). Imaging was performed using a Zeiss Gemini 300 field-emission scanning electron microscope (FE-SEM) from Wisconsin Centers for Nanoscale Technology under standard operating conditions.

### Spike cell orientation analysis

Spike cell orientation was quantified using Fiji/ImageJ (**Fig. 2f, 4e, and S3b, c**). For each experimental condition, three fin rays were analyzed per fish. Within each fin ray, an ROI extending 300 µm along the longitudinal axis was defined, with the width adjusted to the individual fin ray. The orientation of each *avil*⁺ spike was measured relative to the P-to-D axis, with 0° representing the distal direction and 180° representing the proximal direction. Angular measurements were grouped into 20° intervals for visualization. For comparisons of spike cell orientation across regenerative stages (**Fig. 4e**), the proportion of cells oriented within 0–20° was compared with those oriented within 20–180° using Pearson’s chi-square test, followed by two-sided Fisher’s exact tests for pairwise comparisons with Holm correction. ROIs of regenerating fin tissues were restricted to newly regenerated tissue distal to the original amputation plane.

### Spike cell quantification

Spike cell density (**Fig. 4a and S2d**) was quantified in uninjured and regenerating caudal fins of *avil:m-mCherry*. 3^rd^, 4^th^, and 5^th^ fin rays on both the dorsal and ventral sides were analyzed in six zebrafish per time point. In uninjured fins and regenerating fins at 7 and 41 dpa, two consecutive bone segments per fin ray were analyzed. At 1 and 2 dpa, each region of interest (ROI) encompassed whole regenerated tissue within the boundaries of an intra-fin ray, extending from the amputation plane to the distal tip. At 4 dpa, an ROI per intra-fin ray extended distally from the amputation plane for a length equivalent to two consecutive bone segments. Spike cell density was calculated by dividing the number of *avil*⁺ cells by the measured ROI area and normalizing the result to 10,000 µm². Regions with excessive pigmentation that prevented accurate identification and counting of fluorescently labeled cells were not used. Differences in cell density among uninjured and regenerating fins (**Fig. 4a**) were assessed using one-way ANOVA followed by Tukey’s multiple-comparisons test. Intra-fin ray and inter-fin ray cell densities in uninjured fins (**Fig. S2d**) were compared using a two-sided Student’s t-test.

To quantify the number of *avil:m-mCherry⁺* cells and their corresponding morphologies (**Fig. 4c**), 3^rd^, 4^th^, and 5^th^ fin rays on the dorsal side were analyzed in six zebrafish for the uninjured, 1 dpa, and 2 dpa, and in three zebrafish for the 4 dpa group. In uninjured fins, ROIs in the intra-fin ray regions extended 300 µm distally from the midpoint of the fins. At 1 and 2 dpa, each ROI encompassed whole regenerated tissue within the boundaries of an intra-fin ray, extending from the amputation plane to the distal tip. For 4 dpa, two ROIs within the regenerates were analyzed separately: a distal ROI (Dis) encompassing the distal-most 300 µm and a proximal ROI (Pro) extending 300 µm immediately distal to the amputation plane within the proximal redifferentiation zone.

### Cell contact quantification

For quantification of contacts between *avil:m-mCherry*⁺ cells and *mpeg1:EGFP*⁺ macrophages (**Fig. 3b, g, h**), two consecutive bone segments in each of 3^rd^, 4^th^, and 5^th^ fin rays on the dorsal side were analyzed in four zebrafish. ROIs were centered at approximately 50% along the proximodistal axis of each fin ray and encompassed the full width of the fin ray. Contacts between *avil:m-mCherry*⁺ and *mpeg1:EGFP*⁺ cells were scored based on direct physical association. For statistical analysis, each fin ray was treated as the unit of analysis, and the proportions of cell-body and pseudopodial contacts were compared using two-sided exact paired Wilcoxon signed-rank tests.

To compare the frequency of contacts with *mpeg1*⁺ cell between cortical F-actin-enriched epithelial cells and *avil*⁺ spike cells (**Fig. S4**), contacting and total cells in each population were counted within 3^rd^ - 5^th^ fin rays of the dorsal side. For each fish, counts were pooled across the sampled fin rays to calculate the percentage of cells contacting *mpeg1*⁺ cells. The percentages calculated for individual fish were compared between the two cell populations using a two-sided exact Mann–Whitney U test.

### EdU Incorporation Assay

EdU incorporation assays were performed as previously described^105^. Briefly, *avil:lifeact-mNeonGreen* zebrafish were intraperitoneally injected with 10 µL of EdU solution (10 mM in PBS; TCI Chemical, E1057) at 2 dpa. Regenerating fin tissues were collected at the indicated time points and fixed in 4% PFA overnight at 4 °C. Following fixation, fins were washed once with TBS for 10 min and incubated in freshly prepared EdU staining solution containing 1 mM CuSO₄, 50 mM ascorbic acid, 10 mM Alexa Fluor 594 azide (Invitrogen, A10270), and 100 mM Tris buffer, pH 8.0, for 60 min at room temperature. Samples were then washed several times with 0.2% Triton X-100 in TBS and processed for DAPI staining. After incubation with DAPI staining solution, tissues were washed and mounted in Fluoromount-G.

For EdU incorporation quantification (**Fig. 5b, g**), three fin rays were analyzed per fish. ROIs encompassed the entire regenerating region of each fin ray, extending from the original amputation plane to the distal tip. EdU⁺ *avil*⁺ and EdU⁻ *avil*⁺ cells were counted within each ROI, and the percentage of EdU⁺ *avil*⁺ cells was calculated relative to the total number of *avil*⁺ cells. Individual fin-ray measurements were used for data visualization and calculation of mean ± SEM. For statistical comparisons, cell counts from the three fin rays were pooled within each fish to obtain one fish-level percentage per biological replicate. Differences among time points were assessed using the Kruskal–Wallis test followed by Dunn’s multiple-comparisons test with Holm correction.

For morphological classification (**Fig. 5g**), EdU⁺ *avil*⁺ cells were examined using Imaris and classified into three categories based on their morphology and position relative to the epithelial surface: internal spherical cells (Class I), internal cells with apical structures (Class II), and cells bearing an external spike (Class III). The relative proportions of these classes were calculated from pooled cell counts at each time point.

### Hybridization Chain Reaction

*avil:m-mCherry* caudal fins were fixed and sectioned at 16 µm using a cryostat as previously described ^110^. *In situ* hybridization chain reaction (HCR) v3 was carried out as described in ^111^. Buffers were purchased from Molecular Instruments. Briefly, rehydrated samples were incubated in hybridization buffer for 30 min at 37°. The samples were then incubated in probe solution (1 pmol of pooled probes to 250 μl of hybridization buffer) at 37° overnight. Probes were designed using an open-source software ^112^. 33 probe pairs containing the B1 initiator sequence were designed against zebrafish *avil* mRNA (**Table S1)**. The probes were ordered as an oPools Oligo Pool from Integrated DNA Technologies. Following probe hybridization, the samples were washed 4 times for 15 min each with probe wash buffer at 37°, followed by 2 times wash for 5 min with 5x SSCT at room temperature. The samples were incubated in amplification buffer for 30 min at room temperature. Next, samples were incubated in 15 pmol of each snap-cooled hairpin (B1h1 488 and B1h2 488) in 250 μl of amplification buffer at room temperature overnight. Samples were rinsed in 5x SSCT (2 x 5 min, 2 x 30 min, and 1 x 5 min) and immunostaining was performed using anti-Ds-Red (rabbit, 632496, Takara Bio, 1:500) and Alexa Fluor 594 (rabbit, A11037, Life Technologies, 1:500). Cell nuclei were labeled using DAPI (1:4000). Samples were imaged via a Nikon A1Rs laser-scanning Confocal Microscope.

### Drug treatment

For pharmacological inhibition (**Fig. 6j, k and S7**), zebrafish were exposed to Lorlatinib (MCE, HY-12215) by bath immersion. A 12.5mM Lorlatinib stock solution was prepared in 5% DMSO. Following amputation, fish were transferred to 500 mL system water supplemented with 200 µL of the Lorlatinib stock solution to achieve a final concentration of 5 µM Lorlatinib. Fish were maintained at room temperature, and the drug-containing solution was refreshed daily. Control fish were treated with an equivalent volume of vehicle solution containing 5% DMSO. Treatments were carried out from 3 dpa to 5 dpa. Time-lapse imaging was initiated at 3 dpa / 0 dpt and continued following drug treatment at 1 and 2 dpt. As the regenerating area exceeded 300 µm at these stages, the initial ROI was defined as the region extending from the distal edge of the regenerating fin ray toward the amputation plane, spanning the full width of the corresponding fin ray. For subsequent time points, the most proximal pre-existing cell in each fin ray was identified as the proximal reference point, and the ROI extended distally from this cell to the distal edge of the regenerating fin ray. Relative *avil*⁺ cell numbers at 1 and 2 dpt were normalized to the corresponding cell number at 0 dpt for each fish (**Fig. 6k**). These longitudinal data were analyzed using a linear mixed-effects model. Fin regenerate length was measured along the proximodistal axis from the original amputation plane to the distal edge of the regenerating fin at 0, 1, and 2 dpt (**Fig. S7c**). These repeated measurements were analyzed using a linear mixed-effects model. Residual normality was assessed using the Shapiro–Wilk test.

For *avil^+^* cell ablation (**Fig. 7, 8g, S8**), uninjured *avil:BFP-NTRv2* fish were treated with metronidazole (MTZ; Sigma-Aldrich, M3761) by bath immersion. Fish were transferred to system water containing 100 µM MTZ and incubated for 12 h. After treatment, fish were returned to fresh system water for recovery and subsequent imaging. Images were acquired prior to ablation (Pre) and at 12, 36, 60, 84, and 108 hpt, as well as recovery >14 dpt (Reg).

For ablation experiments (**Fig. 7b, c and S8b-e**), we selected the distal-most bifurcation sites of 3^rd^, 4^th^, and 5^th^ fin rays and defined ROI. Each ROI encompassed the two daughter rays and the intervening inter-fin ray region distal to the bifurcation, extending longitudinally for a length equivalent to two bone segments. The same regions were examined longitudinally before Mtz treatment (Pre), at 12, 36, 60, 84, and 108 h post-treatment (hpt), and at the recovery >14dpt (Reg). Within each ROI, *avil:BFP*⁺ cells and *mpeg1:EGFP*⁺ cells were counted acquired using the AxioZoom stereo fluorescence microscope (Zeiss). For **Fig. 7b, c**, *avil*⁺ and *mpeg1*⁺ cell numbers were quantified longitudinally within the same ROIs. Data were obtained from 18 fin-ray ROIs from six fish for Pre and 12, 36, 60, 84, and 108 hpt, and 12 fin-ray ROIs from four fish for Reg. Changes in cell numbers over time were assessed using Friedman repeated-measures tests, followed by paired Wilcoxon signed-rank tests with Holm correction for planned comparisons. For spatial distribution analysis (**Fig. S8c**), intra-fin ray and inter-fin ray cell numbers were compared at each time point using two-sided paired Wilcoxon signed-rank tests with Holm correction for multiple comparisons. For **Fig. S8d and e**, changes in total cell numbers across time points were assessed using Friedman repeated-measures tests, followed by paired Wilcoxon signed-rank tests with Holm correction for planned comparisons.

### Photoconversion and cell tracking

To track *mpeg1*⁺ immune cells over time, photoconversion was performed using transgenic zebrafish expressing the photoconvertible fluorescent protein Dendra2 under the *mpeg1* promoter (*Tg(mpeg1:Dendra2)^uwm12^*). Photoconversion was carried out using an Olympus FV3000 confocal laser scanning microscope. Dendra2-expressing cells within a single bone segment were exposed to 405-nm laser illumination for 5 min to induce photoconversion from green to red fluorescence. To minimize unintended photoconversion during subsequent imaging of the *avil:BFP* reporter, BFP excitation laser power was reduced to 0.2% of the instrument’s maximum laser power setting. No additional photoconversion was detected under these low-power acquisition conditions.

Confocal imaging was acquired over an axial depth of 45 µm. The imaging and quantification area encompassed the photoconverted bone segment and the immediately adjacent distal bone segment along the same fin ray. The same anatomical region was imaged throughout the tracking period, and quantification was restricted to the intra-ray region.

For **Fig. 8d**, *avil*⁺ cells in the adjacent non-conversion segment were classified into three categories based on their physical associations with *mpeg1^+^* cells: non-contacting (NC), contacting non-photoconverted *mpeg1:G* cells (GC), or contacting photoconverted *mpeg1:G->R* cells (RC). The proportion of *avil*⁺ cells in each category was calculated for each fish at the indicated time points. For **Fig. 8e**, photoconverted *mpeg1:G->R* cells identified by the red fluorescence were classified as contacting or non-contacting based on their spatial association with *avil*⁺ spike cells. The proportion of contacting cells was calculated by dividing the number of *mpeg1:G->R* cells contacting *avil*⁺ cells by the total number of *mpeg1:G->R* cells within the two bone segments.

For **Fig. 8f**, the number of photoconverted *mpeg1:G->R* cells at each subsequent time point was normalized to the corresponding baseline count (0 h) for each fish. Differences in normalized retention between control and Mtz-treated groups were assessed using two-sided exact fish-level permutation tests, with Holm correction for multiple comparisons at 12, 36, and 60 h.

### scRNA-seq analysis

For scRNA-seq analysis, we obtained original sequencing files from PRJNA559885 (SRR11886700) and GSE137971 for O1dpa and M-fins, respectively. These profiles were reanalyzed using 10x Genomics cloud service (https://cloud.10xgenomics.com/cloud-analysis). The Danio_rerio.GRCz11 (release 104) version of the zebrafish reference genome and annotation files were downloaded from Ensemble database (ensembl.org). Raw counts were used for scRNA-seq analysis with the Seurat package v5 ^113^. Low quality cells (nUMI ≤ 500, nGene ≤ 250, mitoRatio > 0.2, log_10_GenesPerUMI < 1.0 for O1dpa; and nUMI ≤ 750, nGene ≤ 300, mitoRatio > 0.15, log_10_GenesPerUMI < 1.5 for M-fins) were filtered out. After careful inspection, the 40 principal components (PCs) of the PCA with resolution 1.4 were used for clustering of O1dpa dataset. Regarding M-fins, the 40 PCs of the PCA with resolution 2 were used for clustering. Marker genes to identify cell-types are listed in **Table S2**. For *avil*-enriched epithelial cell subclustering, we first subset *avil*-enriched cluster and then used 40 PCs of the PCA with resolution 2.0 for clustering. In order to determine cell trajectories, RNA velocity was performed by velocyto v0.17.17 ^114^ using the default parameters for single-cell RNA profiles aligned by *cellranger count* (10X Genomics). The resulting loom files were integrated with the Seurat object corresponding to the *avil*-enriched cluster and analyzed using scVelo v0.2.5 in Python^115^. Merged datasets were filtered to retain the top 3,000 genes with at least 20 shared counts between spliced and unspliced transcripts. The data were then processed with PCA and neighborhood graph construction, followed by velocity estimation using the dynamical model. GO-term was analyzed using Metascape ^60^.

To further characterize specialized epithelial populations in an independent dataset, we reanalyzed publicly available zebrafish trunk skin single-nucleus combinatorial indexing RNA-seq (sci-RNA-seq) data from Aman et al. (2023; GEO accession GSE224695). We downloaded the matrix files from the GEO database and subsetted cells annotated as "MLC", "Goblet Cell", "Goblet Cell 2", "Goblet Cell 3", "Ionocyte", "Ionocyte (slc9a3.2+)", and "NCC Ionocyte" clusters. The subsetted data were normalized using SCTransform, followed by Harmony integration. Clustering was performed using 40 PCs at a resolution of 1.2, and the resulting clusters were newly annotated based on their gene expression profiles.

### Quantification and Statistical analysis

Image quantification was performed using Fiji/ImageJ and Imaris (Oxford Instruments). Statistical analyses were performed using R and GraphPad Prism. All statistical tests were two-sided unless otherwise indicated. Data are presented as individual data points with mean ± SEM unless otherwise specified. The numbers of independent fish, fin-ray ROIs, and cells analyzed, together with the definition of individual data points, are provided in the corresponding figure legends and Source Data.

## Data Availability

Published scRNA-seq profiles are available under accession numbers PRJNA559885 (https://trace.ncbi.nlm.nih.gov/Traces/?run=SRR11886700), GSE137971 (https://www.ncbi.nlm.nih.gov/geo/query/acc.cgi?acc=GSE137971), and GSE224695 (https://www.ncbi.nlm.nih.gov/geo/query/acc.cgi?acc=GSE224695). Source data for all figures are provided with this paper.

## Code Availability

The analysis in this study was performed using standard and publicly available R packages as detailed in the Methods section. No custom codes were developed for any analyses in this study. Therefore, no additional code is available for public sharing. The code used for this study is available from the corresponding author upon reasonable request.

## Acknowledgments

We thank the UW–Madison SMPH BRMS staff and members of the Kang Lab for zebrafish care; Rachel Wong and Owen Lawrence for sharing a membrane localization sequence containing plasmid; Addgene and depositors for sharing plasmids; Randall Massey from Wisconsin Centers for Nanoscale Technology for SEM microscopy; Julie Morasch for technical support; and Ken Poss, Kang lab members for comments on the manuscript.

## Author Contributions

Biological Experiments and Analysis: SC, MK, JOH, ARP, YH, TFR, JK Conceptualization and Designing Experiments: SC, MK, JK

Writing, Editing: SC, JK Supervision: AH, JK Funding: AH, JK

## Funding

National Institutes of Health grant R35 GM 137878 (JK) National Institutes of Health grant R01 HL 151522 (JK) National Institutes of Health grant R21 OD 037634 (JK)

National Institutes of Health, University of Wisconsin Carbone Cancer Center Support grant P30 CA 014520 (JK)

Vilas Young Investigator (JK)

National Institutes of Health grant R35 GM 118027 (AH)

## Supplementary Information

**Supplementary Fig. S1 – S9.**

**Supplementary Table S1. Primers used in this study**

**Supplementary Table S2. Marker genes to annotate cell clusters in zebrafish fins**

**Supplementary Movie 1. 3D visualization of *avil*⁺ spike cells in the zebrafish fin epidermis.**

**Supplementary Movie 2. 3D visualization of representative lateral contact sites between *mpeg1*⁺ immune cells and *avil*⁺ spike cells.**

**Supplementary Movie 3. 3D visualization of representative basal contact sites between *mpeg1*⁺ immune cells and *avil*⁺ spike cells.**

**Supplementary Movie 4. 3D visualization of Class I EdU⁺ *avil:l-Green*⁺ cells.**

**Supplementary Movie 5. 3D visualization of Class II EdU⁺ *avil:l-Green*⁺ cells.**

**Supplementary Movie 6. 3D visualization of Class III EdU⁺ *avil:l-Green*⁺ cells.**

## References

1. Sotiropoulou, P. A. & Blanpain, C. Development and homeostasis of the skin epidermis. Cold Spring Harbor perspectives in biology 4, a008383 10.1101/cshperspect.a008383 (2012).

2. Dao, D. P. D. & Le, P. H. Histology, Goblet Cells. In: StatPearls) (2026).

3. Hollenhorst, M. I. & Krasteva-Christ, G. Chemosensory cells in the respiratory tract as crucial regulators of innate immune responses. The Journal of physiology 601, 1555–1572 10.1113/JP282307 (2023).

4. O’Leary, C. E., Schneider, C. & Locksley, R. M. Tuft Cells-Systemically Dispersed Sensory Epithelia Integrating Immune and Neural Circuitry. Annu Rev Immunol 37, 47–72 10.1146/annurev-immunol-042718-041505 (2019).

5. Howitt, M. R., et al. Tuft cells, taste-chemosensory cells, orchestrate parasite type 2 immunity in the gut. Science 351, 1329–1333 10.1126/science.aaf1648 (2016).

6. Brown, T. L., et al. Dermal appendage-dependent patterning of zebrafish atoh1a+ Merkel cells. eLife 12, 10.7554/eLife.85800 (2023).

7. Neubarth, N. L., et al. Meissner corpuscles and their spatially intermingled afferents underlie gentle touch perception. Science 368, 10.1126/science.abb2751 (2020).

8. Smith, R. J. F. Alarm signals in fishes. Reviews in Fish Biology and Fisheries 2, 33–63 10.1007/BF00042916 (1992).

9. Lee, R. T., Asharani, P. V. & Carney, T. J. Basal keratinocytes contribute to all strata of the adult zebrafish epidermis. PloS one 9, e84858 10.1371/journal.pone.0084858 (2014).

10. Peloggia, J., Lush, M. E., Tsai, Y. Y., Wood, C. & Piotrowski, T. Environmental and molecular control of tissue-specific ionocyte differentiation in zebrafish. Development 151, 10.1242/dev.202809 (2024).

11. Yan, J. J. & Hwang, P. P. Novel discoveries in acid-base regulation and osmoregulation: A review of selected hormonal actions in zebrafish and medaka. Gen Comp Endocrinol 277, 20–29 10.1016/j.ygcen.2019.03.007 (2019).

12. Naama, M., et al. Autophagy controls mucus secretion from intestinal goblet cells by alleviating ER stress. Cell Host Microbe 31, 433–446 e434 10.1016/j.chom.2023.01.006 (2023).

13. Le Guellec, D., Morvan-Dubois, G. & Sire, J. Y. Skin development in bony fish with particular emphasis on collagen deposition in the dermis of the zebrafish (Danio rerio). Int J Dev Biol 48, 217–231 10.1387/ijdb.15272388 (2004).

14. Lu, Y. F., et al. Delta/Jagged-mediated Notch signaling induces the differentiation of agr2-positive epidermal mucous cells in zebrafish embryos. PLoS genetics 17, e1009969 10.1371/journal.pgen.1009969 (2021).

15. Rangel-Huerta, E., Guzman, A. & Maldonado, E. The dynamics of epidermal stratification during post-larval development in zebrafish. Developmental dynamics : an official publication of the American Association of Anatomists 250, 175–190 10.1002/dvdy.249 (2021).

16. Guzman, A., Ramos-Balderas, J. L., Carrillo-Rosas, S. & Maldonado, E. A stem cell proliferation burst forms new layers of P63 expressing suprabasal cells during zebrafish postembryonic epidermal development. Biol Open 2, 1179–1186 10.1242/bio.20136023 (2013).

17. Liu, Z., Meng, Y., Ishikura, A. & Kawakami, A. Live tracking of basal stem cells of the epidermis during growth, homeostasis and injury response in zebrafish. Development 151, 10.1242/dev.202315 (2024).

18. Chen, C. H., et al. Multicolor Cell Barcoding Technology for Long-Term Surveillance of Epithelial Regeneration in Zebrafish. Developmental cell 36, 668–680 10.1016/j.devcel.2016.02.017 (2016).

19. Thompson, T. W. & von Moltke, J. Gastrointestinal tuft cells: Recent insights and next questions. Sci Immunol 11, eaea8740 10.1126/sciimmunol.aea8740 (2026).

20. Feng, X., Fluchter, P., De Tenorio, J. C. & Schneider, C. Tuft cells in the intestine, immunity and beyond. Nat Rev Gastroenterol Hepatol 21, 852–868 10.1038/s41575-024-00978-1 (2024).

21. Sato, A. Tuft cells. Anat Sci Int 82, 187–199 10.1111/j.1447-073X.2007.00188.x (2007).

22. Gerbe, F., et al. Intestinal epithelial tuft cells initiate type 2 mucosal immunity to helminth parasites. Nature 529, 226–230 10.1038/nature16527 (2016).

23. von Moltke, J., Ji, M., Liang, H. E. & Locksley, R. M. Tuft-cell-derived IL-25 regulates an intestinal ILC2-epithelial response circuit. Nature 529, 221–225 10.1038/nature16161 (2016).

24. Huang, L., et al. Tuft cells act as regenerative stem cells in the human intestine. Nature 634, 929–935 10.1038/s41586-024-07952-6 (2024).

25. Kotas, M. E., O’Leary, C. E. & Locksley, R. M. Tuft Cells: Context- and Tissue-Specific Programming for a Conserved Cell Lineage. Annu Rev Pathol 18, 311–335 10.1146/annurev-pathol-042320-112212 (2023).

26. Haber, A. L., et al. A single-cell survey of the small intestinal epithelium. Nature 551, 333–339 10.1038/nature24489 (2017).

27. Zhang, W., et al. Neuro-epithelial circuits promote sensory convergence and intestinal immunity. Nature 651, 440–450 10.1038/s41586-025-09921-z (2026).

28. Touhara, K. K., et al. Parasites trigger epithelial cell crosstalk to drive gut-brain signalling. Nature, 10.1038/s41586-026-10281-5 (2026).

29. Stanbery, A. G., et al. Tuft cells promote reactivation of memory Th2 cells and are required for protective immunity to intestinal helminth re-infection. Immunity, 10.1016/j.immuni.2026.02.022 (2026).

30. Ireland, A. S., et al. Basal cell of origin resolves neuroendocrine-tuft lineage plasticity in cancer. Nature 647, 257–267 10.1038/s41586-025-09503-z (2025).

31. Cortez, V. S., et al. IL-25-induced memory type 2 innate lymphoid cells enforce mucosal immunity. Cell 188, 6220–6235 e6222 10.1016/j.cell.2025.08.017 (2025).

32. Ndjim, M., et al. Tuft cell acetylcholine is released into the gut lumen to promote anti-helminth immunity. Immunity 57, 1260–1273 e1267 10.1016/j.immuni.2024.04.018 (2024).

33. Billipp, T. E., et al. Tuft cell-derived acetylcholine promotes epithelial chloride secretion and intestinal helminth clearance. Immunity 57, 1243–1259 e1248 10.1016/j.immuni.2024.03.023 (2024).

34. Yamada, Y., et al. Tuft cell-like carcinomas: novel cancer subsets present in multiple organs sharing a unique gene expression signature. British journal of cancer 127, 1876–1885 10.1038/s41416-022-01957-6 (2022).

35. Chen, Y., et al. Common and specific gene regulatory programs in zebrafish caudal fin regeneration at single-cell resolution. Genome research 35, 202–218 10.1101/gr.279372.124 (2025).

36. Castranova, D., et al. The axillary lymphoid organ is an external, experimentally accessible immune organ in the zebrafish. J Exp Med 222, 10.1084/jem.20241435 (2025).

37. Jones, L. O., et al. Single-cell resolution of the adult zebrafish intestine under conventional conditions and in response to an acute Vibrio cholerae infection. Cell Rep 42, 113407 10.1016/j.celrep.2023.113407 (2023).

38. Willms, R. J., Jones, L. O., Hocking, J. C. & Foley, E. A cell atlas of microbe-responsive processes in the zebrafish intestine. Cell Rep 38, 110311 10.1016/j.celrep.2022.110311 (2022).

39. Wang, W., et al. Changes in regeneration-responsive enhancers shape regenerative capacities in vertebrates. Science 369, 10.1126/science.aaz3090 (2020).

40. Hou, Y., et al. Cellular diversity of the regenerating caudal fin. Sci Adv 6, eaba2084 10.1126/sciadv.aba2084 (2020).

41. Akimenko, M. A., Johnson, S. L., Westerfield, M. & Ekker, M. Differential induction of four msx homeobox genes during fin development and regeneration in zebrafish. Development 121, 347–357 10.1242/dev.121.2.347 (1995).

42. Lewis, V. M., et al. Insulin-like growth factor receptor / mTOR signaling elevates global translation to accelerate zebrafish fin regenerative outgrowth. Developmental biology 502, 1–13 10.1016/j.ydbio.2023.05.008 (2023).

43. Ellett, F., Pase, L., Hayman, J. W., Andrianopoulos, A. & Lieschke, G. J. mpeg1 promoter transgenes direct macrophage-lineage expression in zebrafish. Blood 117, e49–56 10.1182/blood-2010-10-314120 (2011).

44. Walton, E. M., Cronan, M. R., Beerman, R. W. & Tobin, D. M. The Macrophage-Specific Promoter mfap4 Allows Live, Long-Term Analysis of Macrophage Behavior during Mycobacterial Infection in Zebrafish. PloS one 10, e0138949 10.1371/journal.pone.0138949 (2015).

45. Renshaw, S. A., et al. A transgenic zebrafish model of neutrophilic inflammation. Blood 108, 3976–3978 10.1182/blood-2006-05-024075 (2006).

46. Hall, C., Flores, M. V., Storm, T., Crosier, K. & Crosier, P. The zebrafish lysozyme C promoter drives myeloid-specific expression in transgenic fish. BMC Dev Biol 7, 42 10.1186/1471-213X-7-42 (2007).

47. Kelsh, R. N. & Eisen, J. S. The zebrafish colourless gene regulates development of non-ectomesenchymal neural crest derivatives. Development 127, 515–525 10.1242/dev.127.3.515 (2000).

48. Hochgreb-Hagele, T. & Bronner, M. E. A novel FoxD3 gene trap line reveals neural crest precursor movement and a role for FoxD3 in their specification. Developmental biology 374, 1–11 10.1016/j.ydbio.2012.11.035 (2013).

49. Zhao, C., et al. Metaphocytes are IL-22BP-producing cells regulated by ETS transcription factor Spic and essential for zebrafish barrier immunity. Cell Rep 42, 112483 10.1016/j.celrep.2023.112483 (2023).

50. Suratt, B. T., et al. Role of the CXCR4/SDF-1 chemokine axis in circulating neutrophil homeostasis. Blood 104, 565–571 10.1182/blood-2003-10-3638 (2004).

51. Li, L., Yan, B., Shi, Y. Q., Zhang, W. Q. & Wen, Z. L. Live imaging reveals differing roles of macrophages and neutrophils during zebrafish tail fin regeneration. The Journal of biological chemistry 287, 25353–25360 10.1074/jbc.M112.349126 (2012).

52. Kobayashi-Sun, J., et al. Uptake of osteoblast-derived extracellular vesicles promotes the differentiation of osteoclasts in the zebrafish scale. Commun Biol 3, 190 10.1038/s42003-020-0925-1 (2020).

53. Thompson, M. A., et al. The cloche and spadetail genes differentially affect hematopoiesis and vasculogenesis. Developmental biology 197, 248–269 10.1006/dbio.1998.8887 (1998).

54. Larson, J. D., et al. Expression of VE-cadherin in zebrafish embryos: a new tool to evaluate vascular development. Developmental dynamics : an official publication of the American Association of Anatomists 231, 204–213 10.1002/dvdy.20102 (2004).

55. Sumanas, S., Jorniak, T. & Lin, S. Identification of novel vascular endothelial-specific genes by the microarray analysis of the zebrafish cloche mutants. Blood 106, 534–541 10.1182/blood-2004-12-4653 (2005).

56. Verma, A., et al. Endothelial cell-specific chemotaxis receptor (ecscr) promotes angioblast migration during vasculogenesis and enhances VEGF receptor sensitivity. Blood 115, 4614–4622 10.1182/blood-2009-10-248856 (2010).

57. Shin, K., Begeman, I. J., Cao, J. & Kang, J. leptin b and its regeneration enhancer illustrate the regenerative features of zebrafish hearts. Developmental dynamics : an official publication of the American Association of Anatomists 253, 91–106 10.1002/dvdy.556 (2024).

58. Tang, W. J., Watson, C. J., Olmstead, T., Allan, C. H. & Kwon, R. Y. Single-cell resolution of MET- and EMT-like programs in osteoblasts during zebrafish fin regeneration. iScience 25, 103784 10.1016/j.isci.2022.103784 (2022).

59. Koster, M. I., et al. p63 induces key target genes required for epidermal morphogenesis. Proceedings of the National Academy of Sciences of the United States of America 104, 3255–3260 10.1073/pnas.0611376104 (2007).

60. Zhou, Y., et al. Metascape provides a biologist-oriented resource for the analysis of systems-level datasets. Nat Commun 10, 1523 10.1038/s41467-019-09234-6 (2019).

61. Ruppert, A. L., et al. Advillin is a tuft cell marker in the mouse alimentary tract. J Mol Histol 51, 421–435 10.1007/s10735-020-09893-6 (2020).

62. Huang, Y. H., et al. POU2F3 is a master regulator of a tuft cell-like variant of small cell lung cancer. Genes & development 32, 915–928 10.1101/gad.314815.118 (2018).

63. Luo, X. C., et al. Infection by the parasitic helminth Trichinella spiralis activates a Tas2r-mediated signaling pathway in intestinal tuft cells. Proceedings of the National Academy of Sciences of the United States of America 116, 5564–5569 10.1073/pnas.1812901116 (2019).

64. Bannier-Helaouet, M., et al. Human conjunctiva organoids to study ocular surface homeostasis and disease. Cell stem cell 31, 227–243 e212 10.1016/j.stem.2023.12.008 (2024).

65. Ualiyeva, S., et al. A nasal cell atlas reveals heterogeneity of tuft cells and their role in directing olfactory stem cell proliferation. Sci Immunol 9, eabq4341 10.1126/sciimmunol.abq4341 (2024).

66. Gauthier-Kemper, A., et al. Interplay between phosphorylation and palmitoylation mediates plasma membrane targeting and sorting of GAP43. Mol Biol Cell 25, 3284–3299 10.1091/mbc.E13-12-0737 (2014).

67. Choi, J. H., Law, M. Y., Chien, C. B., Link, B. A. & Wong, R. O. In vivo development of dendritic orientation in wild-type and mislocalized retinal ganglion cells. Neural Dev 5, 29 10.1186/1749-8104-5-29 (2010).

68. Hoover, B., et al. The intestinal tuft cell nanostructure in 3D. Sci Rep 7, 1652 10.1038/s41598-017-01520-x (2017).

69. Aman, A. J. & Parichy, D. M. Anatomy, development and regeneration of zebrafish elasmoid scales. Developmental biology 510, 1–7 10.1016/j.ydbio.2024.03.001 (2024).

70. Chang, W. L., et al. The Making of a Flight Feather: Bio-architectural Principles and Adaptation. Cell 179, 1409–1423 e1417 10.1016/j.cell.2019.11.008 (2019).

71. Chen, C. F., et al. Development, regeneration, and evolution of feathers. Annu Rev Anim Biosci 3, 169–195 10.1146/annurev-animal-022513-114127 (2015).

72. Butler, M. T. & Wallingford, J. B. Planar cell polarity in development and disease. Nature reviews Molecular cell biology 18, 375–388 10.1038/nrm.2017.11 (2017).

73. Rodriguez-Boulan, E. & Macara, I. G. Organization and execution of the epithelial polarity programme. Nature reviews Molecular cell biology 15, 225–242 10.1038/nrm3775 (2014).

74. Eom, D. S. & Parichy, D. M. A macrophage relay for long-distance signaling during postembryonic tissue remodeling. Science 355, 1317–1320 10.1126/science.aal2745 (2017).

75. Peterman, E., et al. Zebrafish cutaneous injury models reveal that Langerhans cells engulf axonal debris in adult epidermis. Disease models & mechanisms 16, 10.1242/dmm.049911 (2023).

76. Bowman, R. L., Wang, D. & Eom, D. S. A macrophage subpopulation promotes airineme-mediated intercellular communication in a matrix metalloproteinase-9 dependent manner. Cell Rep 42, 112818 10.1016/j.celrep.2023.112818 (2023).

77. Tan, Y. R., Roan, H. Y. & Chen, C. H. Zebrafish tailfin as an in vivo model for capturing tissue-scale cell dynamics. Seminars in cell & developmental biology 166, 29–35 10.1016/j.semcdb.2024.12.005 (2025).

78. Sehring, I. & Weidinger, G. Zebrafish Fin: Complex Molecular Interactions and Cellular Mechanisms Guiding Regeneration. Cold Spring Harbor perspectives in biology 14, 10.1101/cshperspect.a040758 (2022).

79. Li, Y. H., et al. G protein subunit Ggamma13-mediated signaling pathway is critical to the inflammation resolution and functional recovery of severely injured lungs. eLife 12, 10.7554/eLife.92956 (2024).

80. McGuckin, M. A., Linden, S. K., Sutton, P. & Florin, T. H. Mucin dynamics and enteric pathogens. Nat Rev Microbiol 9, 265–278 10.1038/nrmicro2538 (2011).

81. Asada, R., et al. The endoplasmic reticulum stress transducer OASIS is involved in the terminal differentiation of goblet cells in the large intestine. The Journal of biological chemistry 287, 8144–8153 10.1074/jbc.M111.332593 (2012).

82. Nwako, J. G. & McCauley, H. A. Enteroendocrine cells regulate intestinal homeostasis and epithelial function. Mol Cell Endocrinol 593, 112339 10.1016/j.mce.2024.112339 (2024).

83. Ye, L., et al. Enteroendocrine cells sense bacterial tryptophan catabolites to activate enteric and vagal neuronal pathways. Cell Host Microbe 29, 179–196 e179 10.1016/j.chom.2020.11.011 (2021).

84. Esmaeilniakooshkghazi, A., George, S. P., Biswas, R. & Khurana, S. Mouse intestinal tuft cells express advillin but not villin. Sci Rep 10, 8877 10.1038/s41598-020-65469-0 (2020).

85. Mahmoud, W., et al. Differential expression of villin and advillin by neuroendocrine and tuft cells in the murine lower airways. Cell Tissue Res 402, 1–20 10.1007/s00441-025-04003-y (2025).

86. Huang, H. Anaplastic Lymphoma Kinase (ALK) Receptor Tyrosine Kinase: A Catalytic Receptor with Many Faces. Int J Mol Sci 19, 10.3390/ijms19113448 (2018).

87. Aman, A. J., et al. Transcriptomic profiling of tissue environments critical for post-embryonic patterning and morphogenesis of zebrafish skin. eLife 12, 10.7554/eLife.86670 (2023).

88. Solomon, B. J., et al. Lorlatinib in patients with ALK-positive non-small-cell lung cancer: results from a global phase 2 study. Lancet Oncol 19, 1654–1667 10.1016/S1470-2045(18)30649-1 (2018).

89. Fadeev, A., et al. ALKALs are in vivo ligands for ALK family receptor tyrosine kinases in the neural crest and derived cells. Proceedings of the National Academy of Sciences of the United States of America 115, E630–E638 10.1073/pnas.1719137115 (2018).

90. Guilliams, M. & Scott, C. L. Does niche competition determine the origin of tissue-resident macrophages? Nat Rev Immunol 17, 451–460 10.1038/nri.2017.42 (2017).

91. Park, S., et al. Skin-resident immune cells actively coordinate their distribution with epidermal cells during homeostasis. Nature cell biology 23, 476–484 10.1038/s41556-021-00670-5 (2021).

92. Sharrock, A. V., et al. NTR 2.0: a rationally engineered prodrug-converting enzyme with substantially enhanced efficacy for targeted cell ablation. Nature methods 19, 205–215 10.1038/s41592-021-01364-4 (2022).

93. Barros-Becker, F., Lam, P. Y., Fisher, R. & Huttenlocher, A. Live imaging reveals distinct modes of neutrophil and macrophage migration within interstitial tissues. Journal of cell science 130, 3801–3808 10.1242/jcs.206128 (2017).

94. Peterman, E., et al. Microtubules regulate tissue-level navigation in skin-resident macrophages. Journal of cell science 138, 10.1242/jcs.264101 (2025).

95. Harvie, E. A., Green, J. M., Neely, M. N. & Huttenlocher, A. Innate immune response to Streptococcus iniae infection in zebrafish larvae. Infect Immun 81, 110–121 10.1128/IAI.00642-12 (2013).

96. Buissant des Amorie, J. R., et al. Intestinal tuft cell subtypes represent successive stages of maturation driven by crypt-villus signaling gradients. Nat Commun 16, 6765 10.1038/s41467-025-61878-9 (2025).

97. Kotrschal, K. Solitary chemosensory cells: why do primary aquatic vertebrates need another taste system? Trends in ecology & evolution 11, 110–114 10.1016/0169-5347(96)81088-3 (1996).

98. Osculati, F., et al. The solitary chemosensory cells and the diffuse chemosensory system of the airway. Eur J Histochem 51 **Suppl 1**, 65–72 (2007).

99. Sbarbati, A. & Osculati, F. Solitary chemosensory cells in mammals? Cells Tissues Organs 175, 51–55 10.1159/000073437 (2003).

100. Kotrschal, K., Krautgartner, W. D. & Hansen, A. Ontogeny of the solitary chemosensory cells in the zebrafish, Danio rerio. Chem Senses 22, 111–118 10.1093/chemse/22.2.111 (1997).

101. Kotrschal, K., Royer, S. & Kinnamon, J. C. High-voltage electron microscopy and 3-D reconstruction of solitary chemosensory cells in the anterior dorsal fin of the Gadid fish Ciliata mustela (Teleostei). J Struct Biol 124, 59–69 10.1006/jsbi.1998.4046 (1998).

102. Silverman, J. B., Krystofiak, E. E., Caplan, L. R., Lau, K. S. & Tyska, M. J. Organization of a cytoskeletal superstructure in the apical domain of intestinal tuft cells. The Journal of cell biology 223, 10.1083/jcb.202404070 (2024).

103. Lee, M., et al. Tuft cells shape airway remodeling by eliciting OXGR1- and SOX9-dependent stem cell programs. Nat Commun, 10.1038/s41467-026-70763-y (2026).

104. Nakagawa, A., et al. De novo formation of tuft and goblet cells protects the intrapancreatic biliary duct system from inflammatory injury. FEBS J, 10.1111/febs.70330 (2025).

105. Fuqua, W. C., Winans, S. C. & Greenberg, E. P. Quorum sensing in bacteria: the LuxR-LuxI family of cell density-responsive transcriptional regulators. J Bacteriol 176, 269–275 10.1128/jb.176.2.269-275.1994 (1994).

106. Keating, M., et al. A robust knock-in approach using a minimal promoter and a minicircle. Developmental biology 505, 24–33 10.1016/j.ydbio.2023.10.002 (2024).

107. Varshney, G. K., et al. High-throughput gene targeting and phenotyping in zebrafish using CRISPR/Cas9. Genome research 25, 1030–1042 10.1101/gr.186379.114 (2015).

108. Chertkova, A. O., et al. Robust and Bright Genetically Encoded Fluorescent Markers for Highlighting Structures and Compartments in Mammalian Cells. bioRxiv, 160374 10.1101/160374 (2020).

109. Rodriguez-Parks, A., et al. Advancing knock-in approaches for robust genome editing in zebrafish. Biol Open, 10.1242/bio.062472 (2026).

110. Kang, J., Nachtrab, G. & Poss, K. D. Local Dkk1 crosstalk from breeding ornaments impedes regeneration of injured male zebrafish fins. Developmental cell 27, 19–31 10.1016/j.devcel.2013.08.015 (2013).

111. Choi, H. M., Beck, V. A. & Pierce, N. A. Next-generation in situ hybridization chain reaction: higher gain, lower cost, greater durability. ACS Nano 8, 4284–4294 10.1021/nn405717p (2014).

112. Kuehn, E., et al. Segment number threshold determines juvenile onset of germline cluster expansion in Platynereis dumerilii. J Exp Zool B Mol Dev Evol 338, 225–240 10.1002/jez.b.23100 (2022).

113. Stuart, T., et al. Comprehensive Integration of Single-Cell Data. Cell 177, 1888–1902 e1821 10.1016/j.cell.2019.05.031 (2019).

114. La Manno, G., et al. RNA velocity of single cells. Nature 560, 494–498 10.1038/s41586-018-0414-6 (2018).

115. Bergen, V., Lange, M., Peidli, S., Wolf, F. A. & Theis, F. J. Generalizing RNA velocity to transient cell states through dynamical modeling. Nature biotechnology 38, 1408–1414 10.1038/s41587-020-0591-3 (2020).

