## Supplementary information for "Regenerative epithelial spike cells instruct immune niche homeostasis in zebrafish fin epidermis"

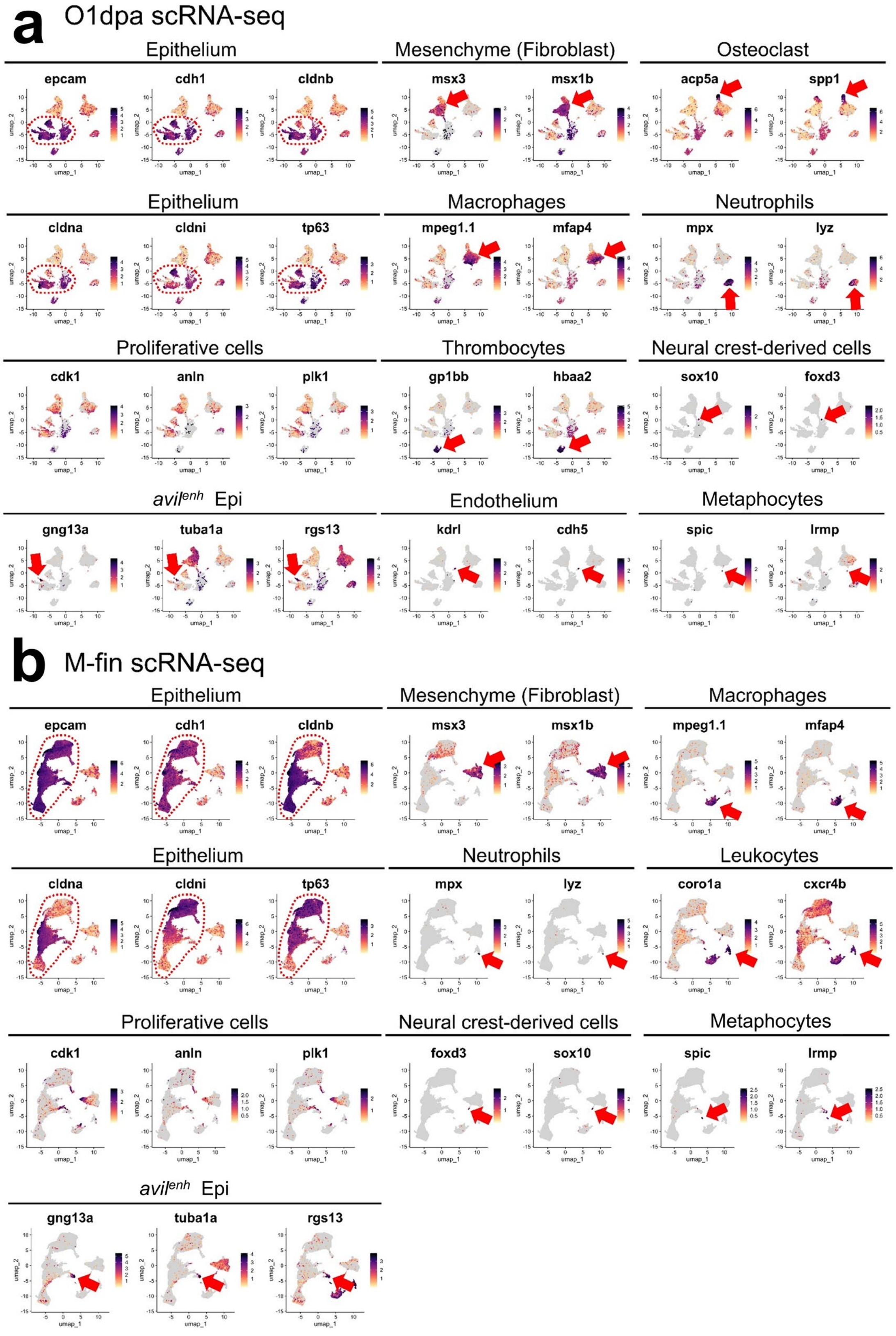

**c**

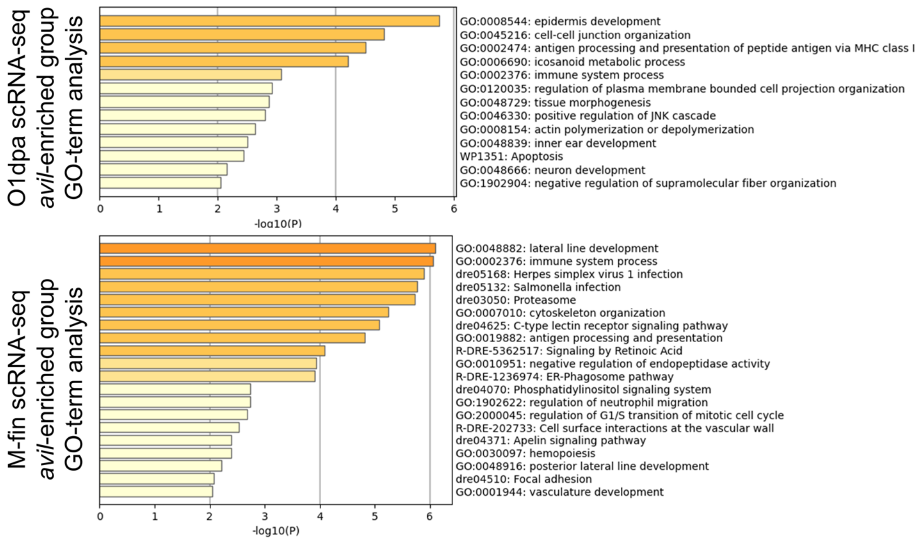

**Supplementary Fig. S1. Marker gene expression and GO-term analysis of *avil*-enriched epithelial populations.** (**a**, **b**) Expressions of marker genes for each subpopulation in O1dpa (**a**) and M-fins (**b**) dataset. The asterisks indicate *avil*-enriched epidermis population. (**c**) GO-term analysis with enriched genes of avil-enriched epitheliual cells

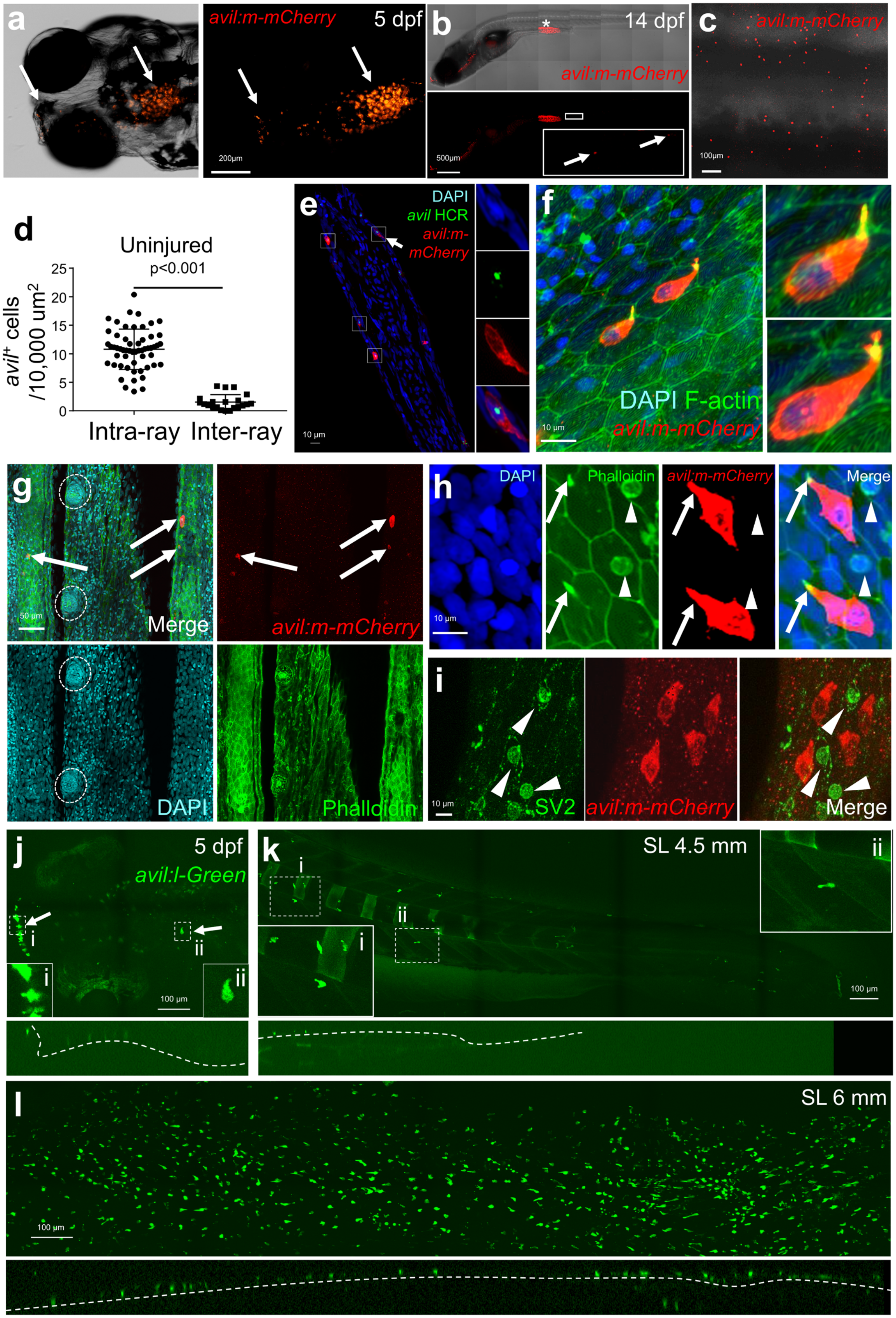

a
avil:m-mCherry
5 dpf
200µm
b
14 dpf
avil:m-mCherry
500µm
c
avil:m-mCherry
100µm
d
Uninjured
p<0.001
avil+ cells /10,000 um2
0
5
10
15
20
25
Intra-ray
Inter-ray
e
DAPI
avil HCR
avil:m-mCherry
10 µm
f
DAPI F-actin
avil:m-mCherry
10 µm
g
50 µm
Merge
avil:m-mCherry
DAPI
Phalloidin
h
DAPI
Phalloidin
avil:m-mCherry
Merge
10 µm
i
SV2
avil:m-mCherry
Merge
10 µm
j
5 dpf
avil:l-Green
i
ii
100 µm
k
SL 4.5 mm
i
ii
100 µm
l
SL 6 mm
100 µm

**Supplementary Fig. S2. Developmental expression and cellular identity of *avil* ⁺ spike cells.** (**a**) Representative images of *avil:m-mCherry* expression in a 5 dpf larva, showing *avil*$^+$ cells in cranial sensory epithelial regions (arrows). (**b**) Representative images of *avil:m-mCherry* expression at 14 dpf. Arrows indicate *avil*$^+$ cells in trunk area. (**c**) Whole-mount image of an adult trunk skin showing wide distribution of *avil:m-mCherry* $^+$ cells. (**d**) Quantification of *avil*$^+$ cell density in intra-ray and inter-ray regions of uninjured fins (n = 56 intra-fin ray and 21 inter-fin ray regions from 6 fish). Statistical significance was assessed using a two-sided Student's t-test. Data are presented with mean ± SD. (**e**) HCR staining showing endogenous *avil* transcript expression in *avil:m-mCherry*$^+$ cells. Enlarged images of the indicated cell are shown in individual fluorescence channels and the merged image. (**f**) Confocal images of uninjured *avil:mem-mCherry* fins stained with Phalloidin and DAPI, showing representative spike cells with prominent apical protrusions enriched in F-actin (arrows). (**g**) Whole-mount confocal images showing the distribution of *avil:m-mCherry*$^+$ cells relative to surrounding neuromasts (dashed circles). Arrows indicate representative *avil*$^+$ spike cells. (**h, i**) Co-labeling analysis showing that *avil:m-mCherry*$^+$ cells are distinct from Merkel cells. Arrows indicate *avil* $^+$ cells, and arrowheads indicate marker-positive SV2$^+$ Merkel cells. (**j–l**) Developmental distribution of *avil*$^+$ cells visualized using the *avil:l-Green* reporter. (**j**) At 5 dpf, *avil:l-Green*$^+$ cells with spike-like protrusions are observed in the cranial sensory epithelial regions. (**k**) At SL 4.5mm, *avil:l-Green*$^+$ cells are sparsely distributed in the trunk epidermis. (**l**) At SL 6 mm, *avil:l-Green*$^+$ cells were observed abundantly in both the trunk epidermis and developing fins. Enlarged regions are shown in the insets. Arrows indicate spike-shaped *avil*$^+$ cells. Scale bars: **a**, 200 µm; **b**, 500 µm; **c**, **j-l**, 100 µm; **e**, **f**, **h**, **i**, 10 µm; **g**, 50 µm.

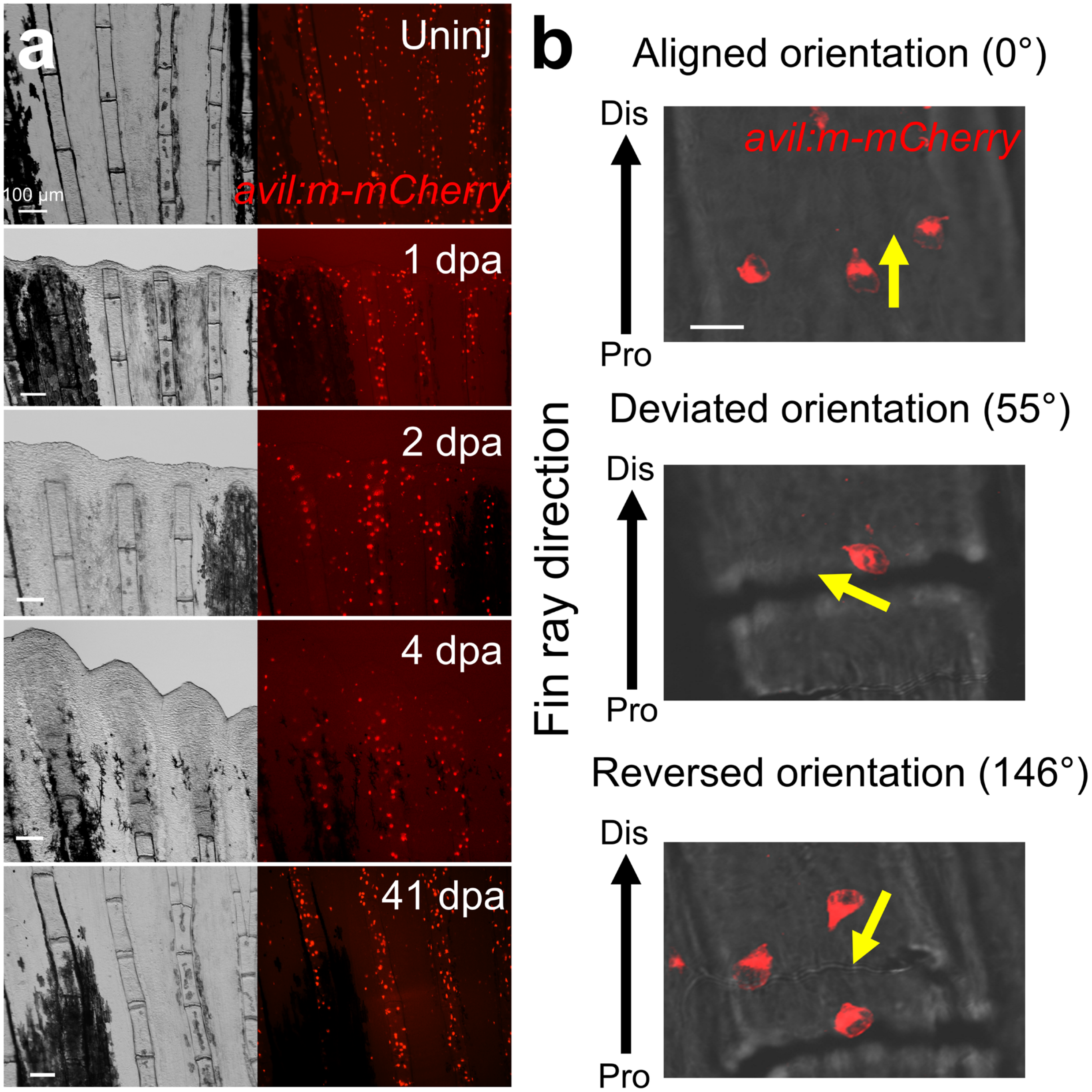

**c**

| Angle | Uninj | 1 dpa | 2 dpa | 4 dpa-Dis | 4 dpa-Pro |
|---|---|---|---|---|---|
| 0º - 20º | 65.89% | 17.65% | 26.81% | 10.65% | 59.77% |
| 20º - 40º | 20.57% | 25.49% | 33.33% | 14.53% | 14.45% |
| 40º - 60º | 7.29% | 17.65% | 15.94% | 13.32% | 7.42% |
| 60º - 80º | 2.08% | 9.80% | 6.52% | 10.17% | 3.13% |
| 80º - 100º | 0.78% | 5.88% | 2.17% | 4.36% | 1.17% |
| 100º - 120º | 0.26% | 11.76% | 5.07% | 7.02% | 0.78% |
| 120º - 140º | 1.30% | 7.84% | 2.17% | 9.69% | 1.95% |
| 140º - 160º | 1.04% | 0.00% | 2.90% | 15.50% | 2.73% |
| 160º - 180º | 0.78% | 3.92% | 5.07% | 14.77% | 8.59% |
| Total cell number | 384 | 51 | 138 | 413 | 256 |

**Supplementary Fig. S3. Dynamic changes in *avil*⁺ spike cell orientation during fin regeneration.** (**a**) Whole-mount bright-field and fluorescence images of *avil:m-mCherry* fins showing the distribution of *avil*$^+$ cells in uninjured fins and at 1, 2, 4, and 41 dpa. (**b**) Representative images illustrating three categories of *avil*$^+$ cell orientation relative to the P-to-D fin ray axis: aligned orientation, deviated orientation, and reversed orientation. Yellow arrows indicate the orientation of individual *avil*$^+$ cells. (**c**) Distribution of *avil*$^+$ spike cell orientation angles grouped into 20° intervals at different regeneration stages. At 4 dpa, Dis and Pro regions were analyzed separately. Total cell counts were 384, 51, 138, 413, and 256 cells for uninjured, 1 dpa, 2 dpa, 4 dpa-Dis, and 4 dpa-Pro, respectively (12 fin rays from 6 fish for uninjured; 9 fin rays from 3 fish for other time points). Scale bar: **a,** 100 μm.

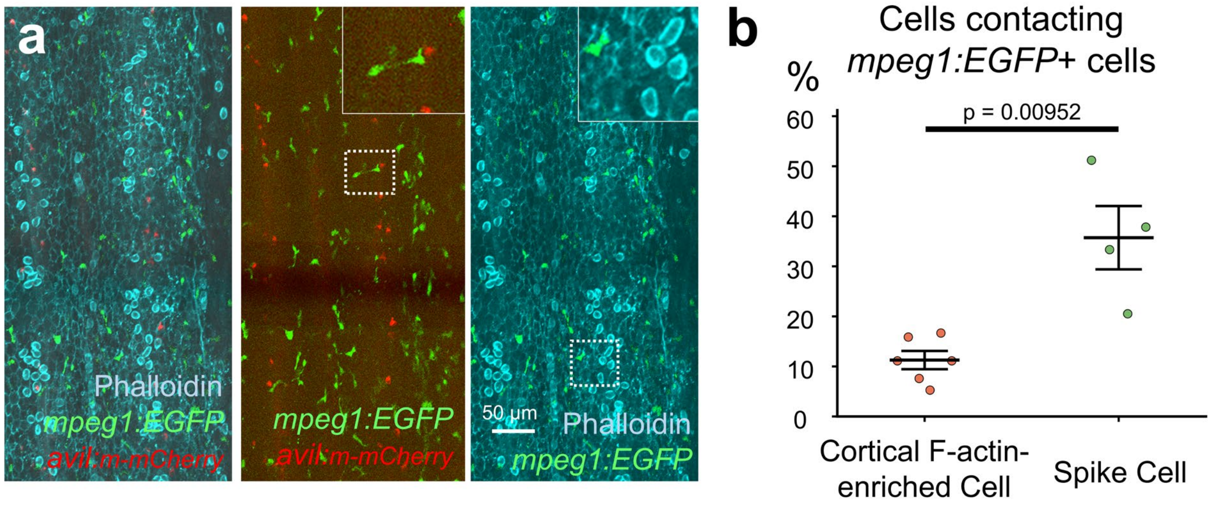

**Supplementary Fig. S4. Comparison of *mpeg1*⁺ immune cell contacts with cortical F-actin-enriched cells and *avil*⁺ spike cells (a)** Representative confocal images showing *mpeg1*⁺ cells (green) in association with *avil*⁺ spike cells (red) and cortical F-actin-enriched epithelial cells visualized by phalloidin staining (cyan). Boxed regions are shown at higher magnification in the insets. **(b)** Quantification of cortical F-actin-enriched epithelial cells and *avil*⁺ spike cells contacting *mpeg1*⁺ cells. Each dot represents one fish (n = 6 and 4 fish for cortical F-actin-enriched epithelial cells and spike cells, respectively). Statistical significance was determined using a two-sided exact Mann–Whitney U test. Data are presented with mean ± SEM. Scale bar: **a,** 50 μm.

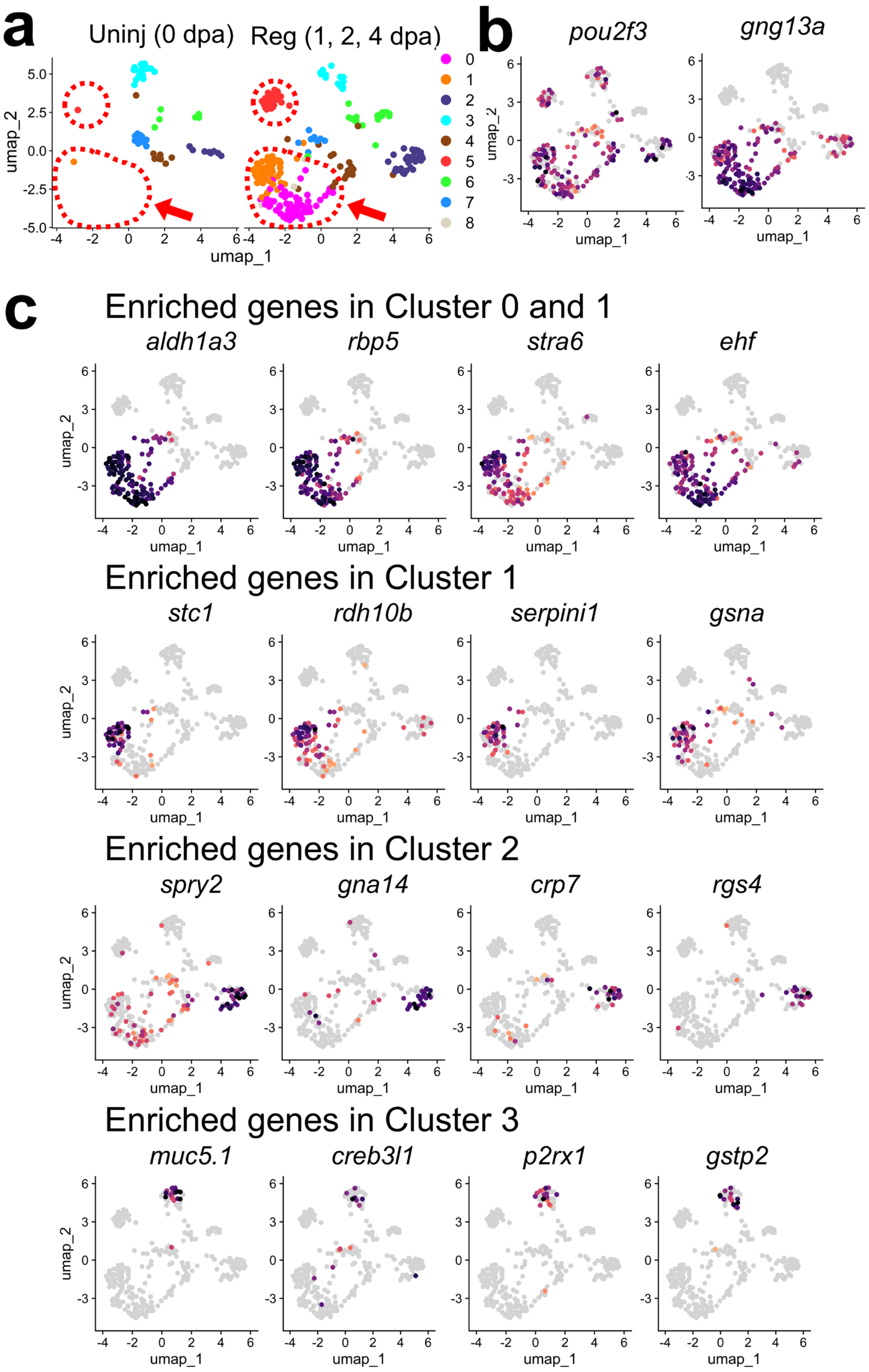

a
Uninj (0 dpa)
Reg (1, 2, 4 dpa)
umap_1
umap_2
b
pou2f3
gng13a
c
Enriched genes in Cluster 0 and 1
aldh1a3
rbp5
stra6
ehf
Enriched genes in Cluster 1
stc1
rdh10b
serpini1
gsna
Enriched genes in Cluster 2
spry2
gna14
crp7
rgs4
Enriched genes in Cluster 3
muc5.1
creb3l1
p2rx1
gstp2

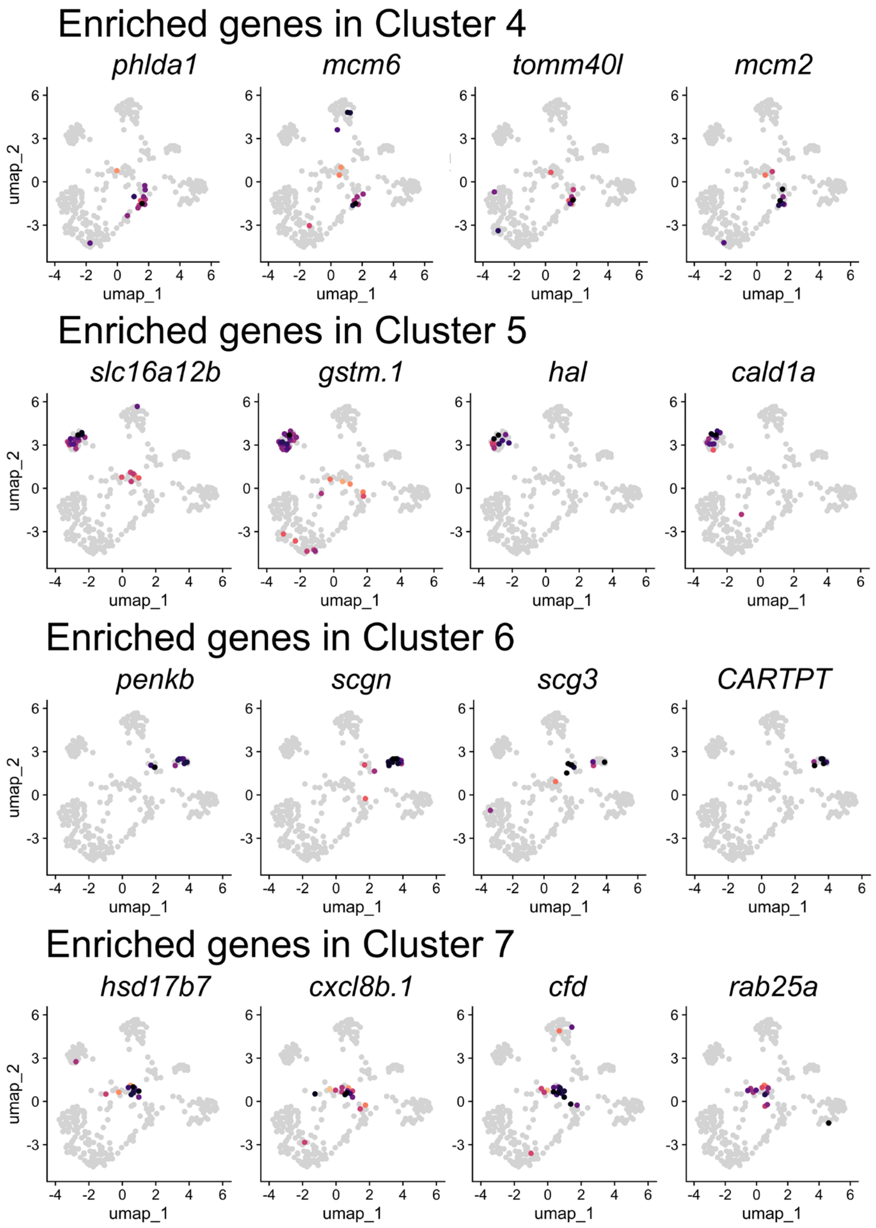

**Supplementary Fig. S5. Molecular heterogeneity of *avil*-enriched epithelial cells in regenerating fins.** (**a**) UMAP visualization of subclusters within the *avil*-enriched epithelial population from uninjured (left) and regenerating (right) fins. Regeneration-emerging cell clusters are highlighted by red dotted circles. (**b**) Featureplots showing *pou2f3* and *gng13a* for *avil*-enriched epithelial subcluster. (**c**) Expressions of marker genes for each cluster in *avil*-enriched epidermis population

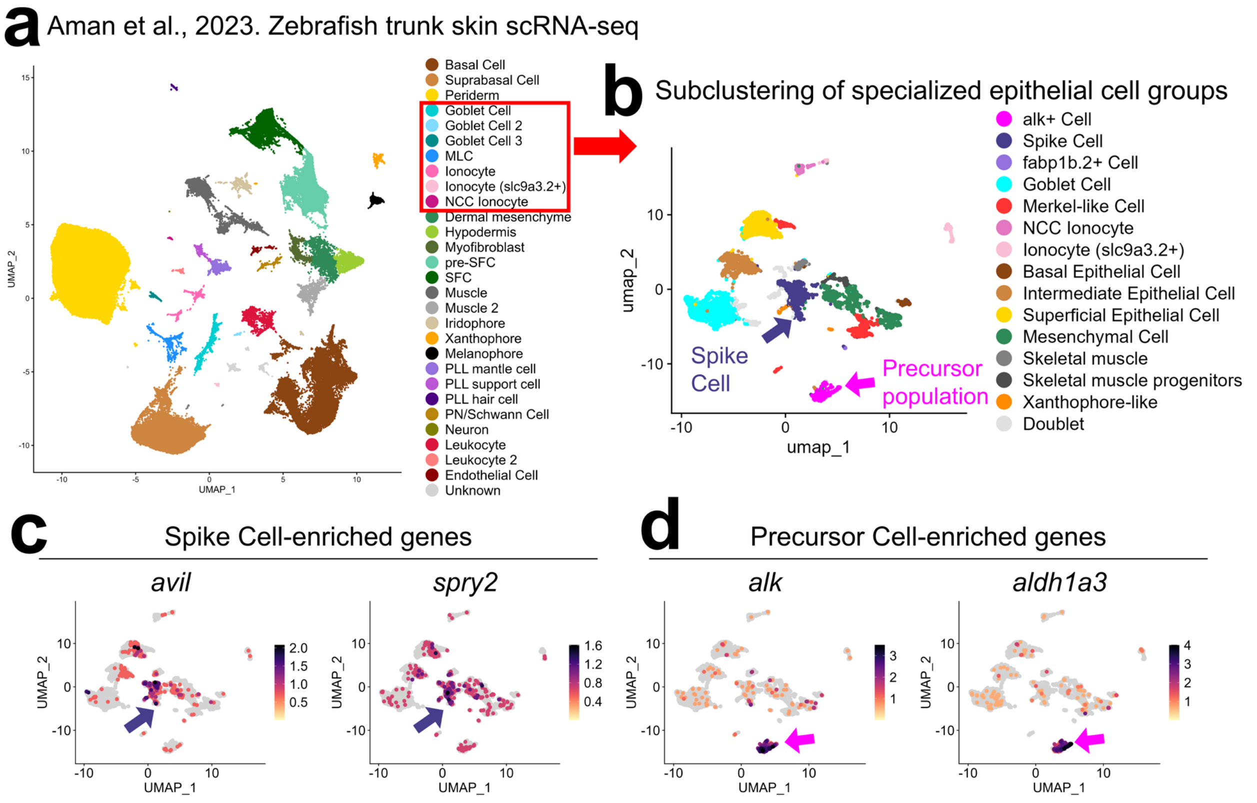

**Supplementary Fig. S6. Zebrafish trunk skin scRNA-seq analysis identifies cell clusters equivalent to spike cell and their precursor populations. (a)** UMAP visualization of the published zebrafish trunk skin scRNA-seq dataset from Aman et al. (2023). The red box highlights the specialized epithelial cell populations selected for further subclustering analysis. **(b)** UMAP visualization of the subclustered epithelial populations, including a spike cell-like cluster and a putative precursor population enriched for *alk* expression (arrows). **(c)** Feature plots showing the expression of *avil* and *spry2* (purple arrows), which are enriched in the spike cell population. **(d)** Feature plots showing the expression of *alk* and *aldh1a3* (magenta arrows), which are enriched in the precursor population.

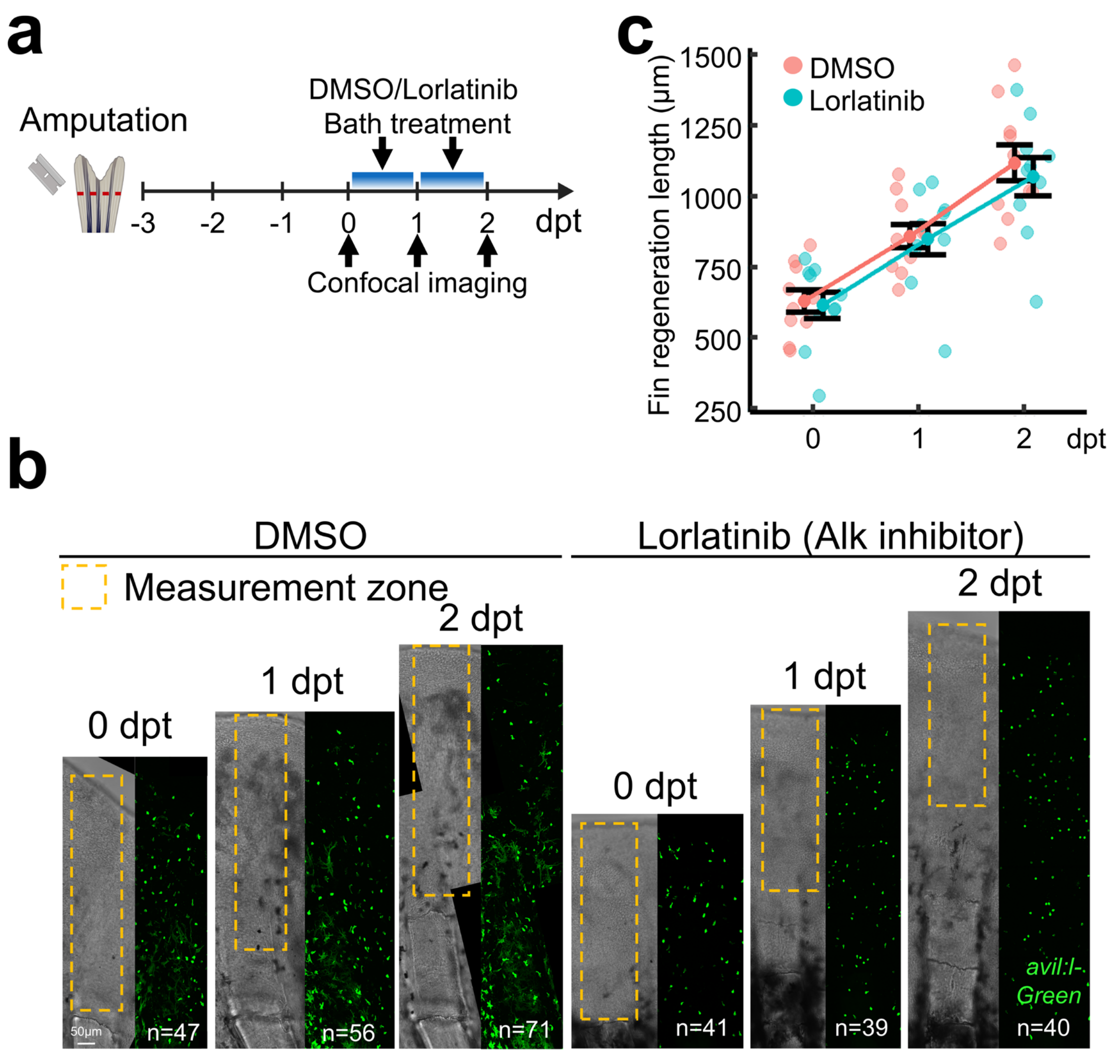

**Supplementary Fig. S7. Lorlatinib, Alk inhibitor, treated fins.** (**a**) Experimental scheme illustrating fin amputation, followed by DMSO or Lorlatinib water bath treatment and longitudinal confocal imaging at 0, 1, and 2 dpt. (**b**) Representative whole-mount confocal images showing the distribution of *avil*$^{+}$ spike cells in regenerating fins treated with DMSO or Lorlatinib at 0, 1, and 2 dpt. Dashed boxes indicate the measurement zones used for **Fig. 6j**. (**c**) Quantification of DMSO- and Lorlatinib-treated fin regenerate length at 0, 1, and 2 dpt. Each dot represents an individual fish with the same fish measured repeatedly across time points. Data are presented with mean ± SEM. Scale bar: **B**, 50 μm.

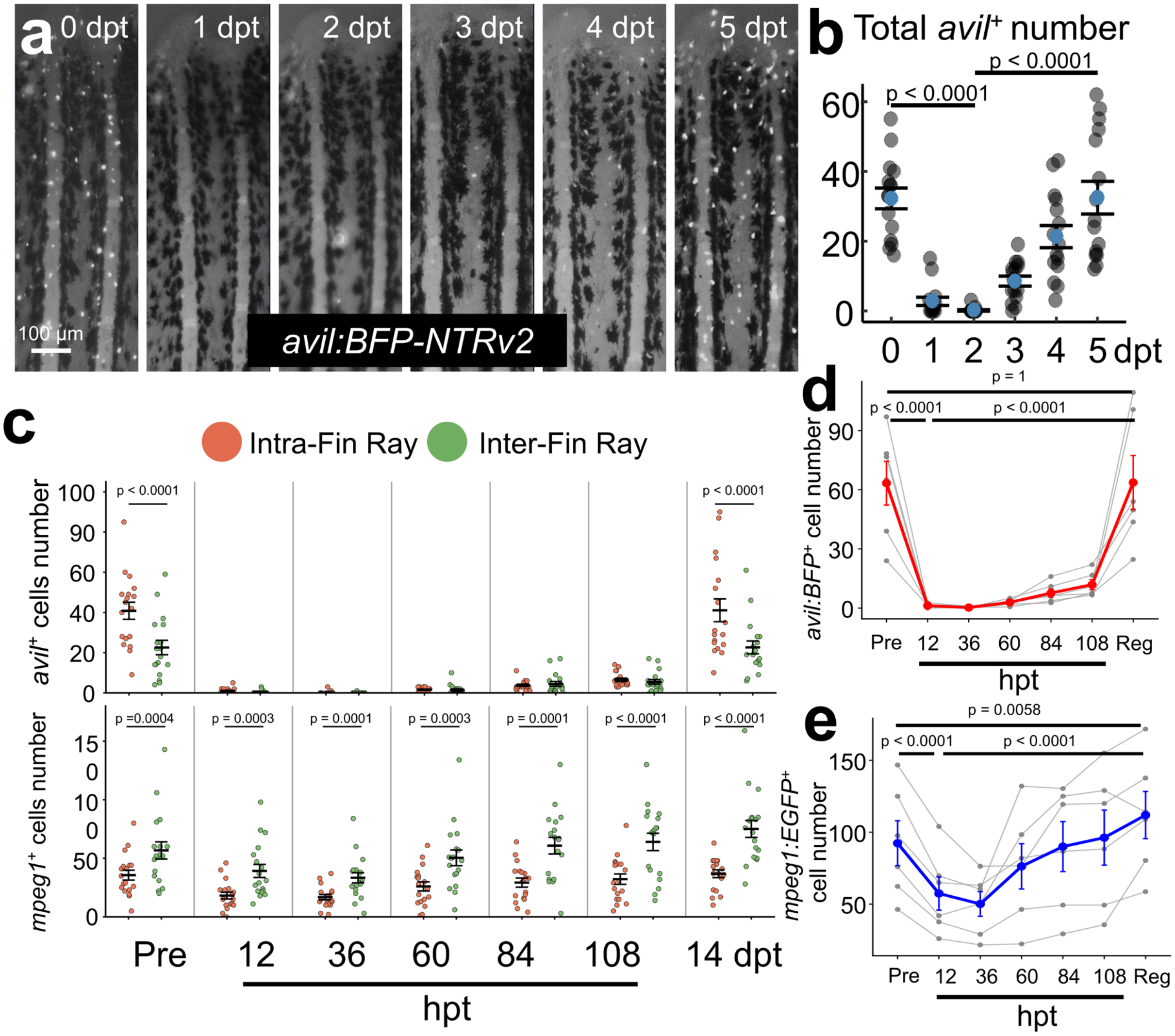

**Supplementary Fig. S8. Mtz-induced ablation and recovery of *avil*⁺ spike cells.** (**a**) Representative whole-mount images of *avil*:BFP-NTRv2 fins from 0 to 5 dpt, showing rapid depletion of *avil*⁺ cells after ablation and their subsequent recovery over time. (**b**) Quantification of *avil*⁺ cell numbers from 0 to 5 dpt. Each dot represents an individual fin ray (n = 15 fin rays from 5 fish), with repeated measurements across time points. Statistical significance was assessed using the Friedman test followed by paired Wilcoxon signed-rank tests with Holm correction. (**c**) Quantification of *avil*⁺ spike cells (top) and *mpeg1*$^{+}$ immune cells (bottom) in intra-fin ray and inter-fin ray regions before treatment (Pre), at 12, 36, 60, 84, and 108 hpt, and at 14 dpt. Each dot represents one fin ray ROI (n = 18 fin rays from 6 fish). Intra-fin ray and inter-fin ray counts were compared using two-sided paired Wilcoxon signed-rank tests with Holm correction. (**d**, **e**) Longitudinal quantification of total *avil*$^{+}$ spike cell numbers (**d**) and *mpeg1*$^{+}$ immune cell numbers (**e**) across the indicated time points. Each gray line represents an individual fish, with measurements averaged across three fin-ray ROIs per fish (n = 6 fish; 18 ROIs in total). Adjusted P values are indicated in the panels. Data are presented with mean ± SEM. Scale bar: **a,** 100 µm.

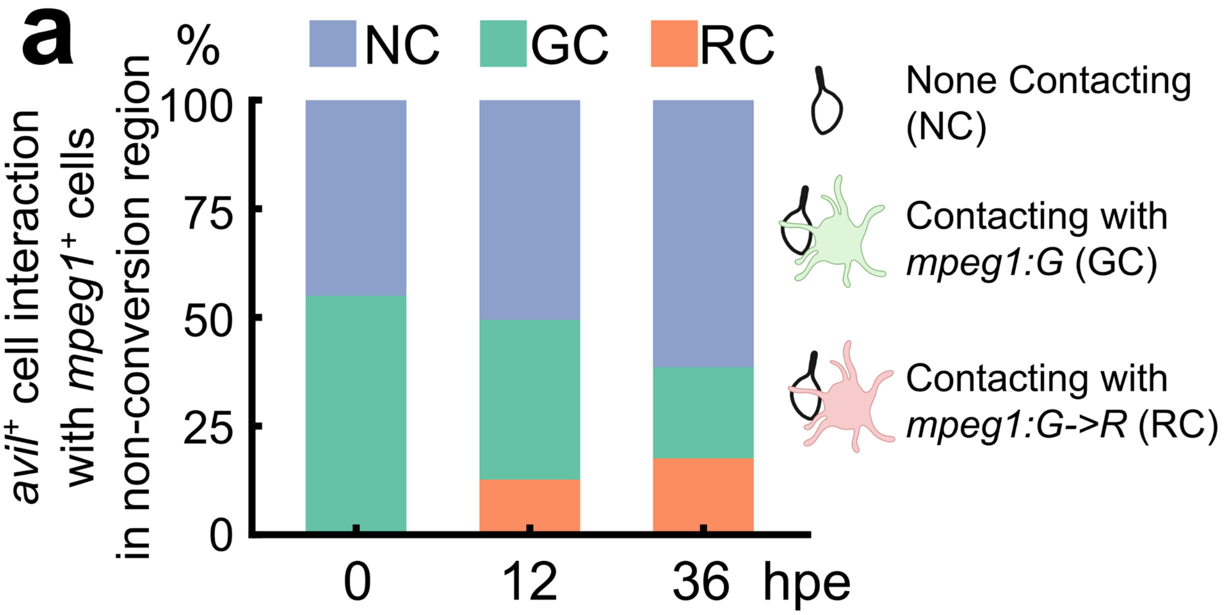

**Supplementary Fig. S9. Dynamic changes of *avil*⁺ spike cell interaction status with *mpeg1*⁺ immune cells in the non-conversion zone.** (**a**) Quantification of the proportions of *avil*⁺ cells classified as non-contacting (NC), contacting with non-photoconverted green *mpeg1:G* cells (GC), or contacting with photoconverted red *mpeg1:G->R* cells (RC) at 0, 12, and 36 hpe. Quantification was performed using non-conversion zone (the fin-ray segment adjacent to the photoconverted segment). Percentages were calculated relative to the total number of *avil*⁺ cells within the same region for each fish. Bars represent mean percentages across fish (n = 4 fish).

### Supplementary Table S1. Primers used in this study

| *avil:memb-mCherry* | |
|---|---|
| avil 5' HA SpeI -f | gg actagt TGGATGTTTGGACTGGGAAC |
| avil 5' HA NheI -r | cga gctagc GATTCAGCGGCGTGTGCGAGA |
| avil 3' HA ClaI -f | aggc atcgat GTGGTGCTTCTGGGATTTAACACA |
| avil 3' HA BamHi -r | act ggatcc TGGAATGATCTCCTCGAACC |
| mem-kozak-NheI -f | cga gctagc GCCACC ATGCTGTGCTGCATCAGA |
| mem HindIII -r | ggt aagctt GGTGGCGACCGGTgggtcc CGGGCCCGCGGTACCGT |

| *avil:lifeact-mNeonGreen* | |
|---|---|
| avil 5'HA-CAGT BsaI -f | atcggtctctCAGTCCTCTAAATCTGCTGTATCTCCT |
| avil 5'HA AGTC BsaI -r | gatGGTCTCTGACTGATTCAGCGGCGTGTGCGAGA |
| Lifeact-BsaI-AGTC-f | atcggtctctAGTCgccaccatgggcgtggccgacctgat |
| SV40-pA-BsaI-CTCA-r | gatggtctctTGAGtaagatacattgatgagtttggaca |
| avil 3'HA CTCA BsaI -f | atcGGTCTCTCTCAGTGGTGCTTCTGGGATTTAACACA |
| avil 3'HA CCAA BsaI -r | gatGGTCTCTTTGGTGGAATGATCTCCTCGAACC |

| *avil:BFP-NTRv2* | |
|---|---|
| pMC-ME-mCherry-NTRv2 | |
| kozak-EGFP-BsaI-AGTC-f | atcggtctctAGTCgccaccatggtgagcaagggcgagga |
| SV40-pA-BsaI-CTCA-r | gatggtctctTGAGgaattaaaaaacctcccacacct |
| pMC-ME-BFP-NTRv2 | |
| BFP-P2A-NTR-vec-f | CAAACTGGGGCACAAGCTTAATggaagcggagctactaacttcagcc |
| BFP-LS-vec-r | TAATCAGCTCGCTCATggtggcGACTagagaccggg |
| BFP-LS-ins-f | cccggtctctAGTCgccaccATGAGCGAGCTGATTAAGGAGA |
| BFP BsaI mut -r | TGCTGCTCGACGTAcGTCTCGTTGTTGGCCTCCTTGA |
| BFP BsaI Mut -f | CAACGAGACgTACGTCGAGCAGCACGAGGT |
| BFP-P2A-NTR-ins-r | tgaagttagtagctccgcttccATTAAGCTTGTGCCCCAGTTTG |

*avil* HCR probe

| Pool name | Sequence |
|---|---|
| B1_avil_33_Dla100 | GAGGAGGGCAGCAAACGGaaTGATGAACATCTCCATCTGCCTATA |
| B1_avil_33_Dla100 | ACTCCTCATTGAATTAGTGAGAGTAtaGAAGAGTCTTCCTTTACG |
| B1_avil_33_Dla100 | GAGGAGGGCAGCAAACGGaaCGAACACTGCCTGGAAATCTGCCTC |
| B1_avil_33_Dla100 | AACAGCTGAACGCCTCCTTAGAGATtaGAAGAGTCTTCCTTTACG |
| B1_avil_33_Dla100 | GAGGAGGGCAGCAAACGGaaTCAGAGATGATGTTATTCTCTGGAG |
| B1_avil_33_Dla100 | TTGTTCACCAGCAGCGCCGCTGGATtaGAAGAGTCTTCCTTTACG |
| B1_avil_33_Dla100 | GAGGAGGGCAGCAAACGGaaCTTGCTAGGGTCCCAGGCAGTGAAC |
| B1_avil_33_Dla100 | CTGCTCGTACGTTTTTCCTCCACTCtaGAAGAGTCTTCCTTTACG |
| B1_avil_33_Dla100 | GAGGAGGGCAGCAAACGGaaGGTACTCGGCACAGGTAGGCACACA |

| | |
|---|---|
| B1_avil_33_Dla100 | GGTCTCTGGAGCCGGGGTGTGTCCGtaGAAGAGTCTTCCTTTACG |
| B1_avil_33_Dla100 | GAGGAGGGCAGCAAACGGaaCGCAGGTCGTCCTGTGTGAACTGCG |
| B1_avil_33_Dla100 | GTGTCCAGCAGCATGACGTCGTCCTtaGAAGAGTCTTCCTTTACG |
| B1_avil_33_Dla100 | GAGGAGGGCAGCAAACGGaaTCGGTCGCTGGCGTATGGCGTTCGT |
| B1_avil_33_Dla100 | GTGCTCCAGAGTCACCTGCTGCAAAtaGAAGAGTCTTCCTTTACG |
| B1_avil_33_Dla100 | GAGGAGGGCAGCAAACGGaaTCCTGACCCTCCGCCATCACCTGCT |
| B1_avil_33_Dla100 | CCCAGACTCTGCCAGAACTCCTCCGtaGAAGAGTCTTCCTTTACG |
| B1_avil_33_Dla100 | GAGGAGGGCAGCAAACGGaaTCGCCATCGCACGCTCGTCCCCGCT |
| B1_avil_33_Dla100 | AGCGACCCATGATCACACTCAGATCtaGAAGAGTCTTCCTTTACG |
| B1_avil_33_Dla100 | GAGGAGGGCAGCAAACGGaaGCTCTGGCTCTTCAGCAGGAACACG |
| B1_avil_33_Dla100 | GCCTTTCCCGTACCACAGGTAAACTtaGAAGAGTCTTCCTTTACG |
| B1_avil_33_Dla100 | GAGGAGGGCAGCAAACGGaaGGAACCTCCACGGCGCGAGTGCTGG |
| B1_avil_33_Dla100 | TTGGAGTTGAGCGACGCCGCCAGCGtaGAAGAGTCTTCCTTTACG |
| B1_avil_33_Dla100 | GAGGAGGGCAGCAAACGGaaGGACGGGCGGCTCTGGCTCCACTGG |
| B1_avil_33_Dla100 | GGTGCGAGCCGCAGATCTGGAACAGtaGAAGAGTCTTCCTTTACG |
| B1_avil_33_Dla100 | GAGGAGGGCAGCAAACGGaaCACCATCTTGCCCTTGAAGATGGAG |
| B1_avil_33_Dla100 | CTTACGGGACGTCCCACCCTCATAGtaGAAGAGTCTTCCTTTACG |
| B1_avil_33_Dla100 | GAGGAGGGCAGCAAACGGaaACGGACACCTGCACCGGCTGACCTC |
| B1_avil_33_Dla100 | AAGTGTCGTGGCTCCTTCCCCATGGtaGAAGAGTCTTCCTTTACG |
| B1_avil_33_Dla100 | GAGGAGGGCAGCAAACGGaaACGCACACGCCGTCACTTCATCCTG |
| B1_avil_33_Dla100 | ACTGCTGATCCACCGTCACCGCATGtaGAAGAGTCTTCCTTTACG |
| B1_avil_33_Dla100 | GAGGAGGGCAGCAAACGGaaCCTCCATAGAAAAACCCGTGTGTAG |
| B1_avil_33_Dla100 | TAGCTGTAGAGAATCAGATAGCAGTtaGAAGAGTCTTCCTTTACG |
| B1_avil_33_Dla100 | GAGGAGGGCAGCAAACGGaaCGCCACCTCCGGCATCATGTGCATC |
| B1_avil_33_Dla100 | GCCGTCGTCCACCATGCGCTCCTGCtaGAAGAGTCTTCCTTTACG |
| B1_avil_33_Dla100 | GAGGAGGGCAGCAAACGGaaTCAGCGATTTTTCCAACAGTGTGTG |
| B1_avil_33_Dla100 | CTGGCATCAAACTTCTCCTGAGGAAtaGAAGAGTCTTCCTTTACG |
| B1_avil_33_Dla100 | GAGGAGGGCAGCAAACGGaaCTGTCCAGTTCTTGAAGAGCTGCTT |
| B1_avil_33_Dla100 | GGCCCAGGCCCTGCGTCTGGTCTTTtaGAAGAGTCTTCCTTTACG |
| B1_avil_33_Dla100 | GAGGAGGGCAGCAAACGGaaGGACTCCACTTTGGTGCTGAGCGGG |
| B1_avil_33_Dla100 | CAGAGCTGATTCTCCTCCGTCACAGtaGAAGAGTCTTCCTTTACG |
| B1_avil_33_Dla100 | GAGGAGGGCAGCAAACGGaaCGGGTCATCGCTGCTTGTCTCTCCG |
| B1_avil_33_Dla100 | TTCTTCAGCTTGATGAACTCCAGAGtaGAAGAGTCTTCCTTTACG |
| B1_avil_33_Dla100 | GAGGAGGGCAGCAAACGGaaCATGATTCAGCAGGTCCTGGGTTAG |
| B1_avil_33_Dla100 | CCCCCTGGTCCAAGATGTAGCAGTCtaGAAGAGTCTTCCTTTACG |
| B1_avil_33_Dla100 | GAGGAGGGCAGCAAACGGaaCGTCCCGTTGGCATCTGACACATGG |
| B1_avil_33_Dla100 | ACTGGTTGCGATCTCGGTGATCTGCtaGAAGAGTCTTCCTTTACG |
| B1_avil_33_Dla100 | GAGGAGGGCAGCAAACGGaaTCAGTGACGTCGTCCGGCGTGCCGC |
| B1_avil_33_Dla100 | AGGGTGAGTTTGGCCATCTGCTCGCtaGAAGAGTCTTCCTTTACG |
| B1_avil_33_Dla100 | GAGGAGGGCAGCAAACGGaaTGCTCAGCATCACCTGCATCAGCAG |
| B1_avil_33_Dla100 | GTAGGGTCGAGGGTCGCTCTCCCAGtaGAAGAGTCTTCCTTTACG |
| B1_avil_33_Dla100 | GAGGAGGGCAGCAAACGGaaCACCCCGATCTCTGCCCGCCCGCCG |
| B1_avil_33_Dla100 | AGCGGCGGCCTCCGCATCGCCCTCAtaGAAGAGTCTTCCTTTACG |

| | |
|---|---|
| B1_avil_33_Dla100 | GAGGAGGGCAGCAAACGGaaTGGTCTTTCCCAGATCCAGCAGAAA |
| B1_avil_33_Dla100 | TGCTTTCCGGCCCGTTCCACTGGATtaGAAGAGTCTTCCTTTACG |
| B1_avil_33_Dla100 | GAGGAGGGCAGCAAACGGaaTGGATGTTATAAGTGTTGGTCTCCA |
| B1_avil_33_Dla100 | CTGCGTCCCTTCACGTGCAGCAGTCtaGAAGAGTCTTCCTTTACG |
| B1_avil_33_Dla100 | GAGGAGGGCAGCAAACGGaaACTTATAGATGATGCCCTGTTTGAA |
| B1_avil_33_Dla100 | GGTTCATCCCTGAAGCCACTCCACCtaGAAGAGTCTTCCTTTACG |
| B1_avil_33_Dla100 | GAGGAGGGCAGCAAACGGaaCTGGACCTCGCGGTGCTGCACCGGT |
| B1_avil_33_Dla100 | TCCGCAGAACATGCTGGACTCGTGCtaGAAGAGTCTTCCTTTACG |
| B1_avil_33_Dla100 | GAGGAGGGCAGCAAACGGaaGCGTACATCGCTGCGGCGCCCTGCT |
| B1_avil_33_Dla100 | GAGCCCAGGAACTCGTCCAGCTGGAtaGAAGAGTCTTCCTTTACG |
| B1_avil_33_Dla100 | GAGGAGGGCAGCAAACGGaaAATGGATGTCATAAGATATCGTTTT |
| B1_avil_33_Dla100 | CTATTGACGATGCAGAGCCGATCCAtaGAAGAGTCTTCCTTTACG |
| B1_avil_33_Dla100 | GAGGAGGGCAGCAAACGGaaATCAAAACCAGATCCATCTTCTCGA |
| B1_avil_33_Dla100 | AAAAACTGTCCATGAAACTTTTCTGtaGAAGAGTCTTCCTTTACG |

**Supplementary Table S2. Marker genes to annotate cell clusters in zebrafish fins**

| Description | Markers |
|---|---|
| Epithelium | |
| *avil*-enriched epithelium | *epcam, cdh1, cldnh, pou2f3,agr2, avil, gng13a* |
| Superficial epithelium | *epcam, cdh1, cldnb, cldne, cldnf* |
| Intermediate epithelium | *epcam, cdh1, tp63, cldna, cldnb* |
| Basal epithelium | *epcam, cdh1, tp63, cldni, col7a1, col17a1a, col17a1b* |
| Reg-specific Basal epithelium | *epcam, cdh1, tp63, cldni, col7a1, col17a1b* |
| Proliferative epithelium | *epcam, cdh1, tp63, cdk1, anln, plk1, rrm2.1* |
| Mesenchyme/fibroblast | |
| lepb enriched Mes blastema | *msx1b, msx3, twist1a, lepb, tph1b* |
| lepb low Mes blastema | *msx1b, msx3, twist1a* |
| Osteoblasts | *msx1b, msx3, twist1a, sp7, bmp2b, spp1, bgna* |
| Mes-uninj | *msx1b, msx3, twist1a* |
| Differentiated fibroblasts | *msx1b, msx3, twist1a, fhl1a, olfml2ba* |
| Immune cells | |
| Macrophages | *mpeg1.1, mfap4* |
| Neutrophils | *mpx, lyz, mmp13a* |
| Thrombocytes | *gp1bb, gp9, hbaa2* |
| Metaphocytes | *epcam, cdh1, mpeg1.1, spic, lrmp* |
| Leukocytes | *coro1a, cxcr4b* |
| Others | |
| Osteoclasts | *acp5a (trap), spp1* |
| Neural crest derived cells | *sox10, foxd3, sox2, bco2b, bscl2l, gch2* |
| Endothelial cells | *kdrl, ecscr, cdh5* |

**Supplementary Movie 1. 3D visualization of *avil*$^{+}$ spike cells in the zebrafish fin epidermis.**

**Supplementary Movie 2. 3D visualization of representative lateral contact sites between *mpeg1*$^{+}$ immune cells and *avil*$^{+}$ spike cells.**

**Supplementary Movie 3. 3D visualization of representative basal contact sites between *mpeg1*$^{+}$ immune cells and *avil*$^{+}$ spike cells.**

**Supplementary Movie 4. 3D visualization of Class I EdU$^{+}$ *avil:l-Green*$^{+}$ cells.**

**Supplementary Movie 5. 3D visualization of Class II EdU$^{+}$ *avil:l-Green*$^{+}$ cells.**

**Supplementary Movie 6. 3D visualization of Class III EdU$^{+}$ *avil:l-Green*$^{+}$ cells.**
